# PlexinA4-Semaphorin3A mediated crosstalk between main cortical interneuron classes is required for superficial interneurons lamination

**DOI:** 10.1101/2020.06.23.166926

**Authors:** Greta Limoni, Mathieu Niquille, Sahana Murthy, Denis Jabaudon, Alexandre Dayer

## Abstract

In the mammalian cerebral cortex, the developmental events governing the allocation of different classes of inhibitory interneurons (INs) into distinct cortical layers are poorly understood. Here we report that the guidance receptor PlexinA4 (PLXNA4) is upregulated in serotonin receptor 3a-expressing (HTR3A^+^) cortical INs (hINs) as they invade the cortical plate and that it regulates their laminar allocation to superficial cortical layers. We find that the PLXNA4 ligand Semaphorin3A (SEMA3A) acts as a chemorepulsive factor on hINs migrating into the nascent cortex and demonstrate that SEMA3A specifically controls their laminar positioning through PLXNA4. We identify that deep layer INs constitute a major source of SEMA3A in the developing cortex and demonstrate that cell-type specific genetic deletion of SEMA3A in these INs specifically affects the laminar allocation of hINs. These data demonstrate that in the neocortex, deep layer INs control the laminar allocation of hINs into superficial layers.

## Introduction

GABAergic interneurons (INs) are a highly heterogeneous population that plays critical roles in cortical circuits by controlling the activity of glutamatergic excitatory neurons (Fishell and Kepecs 2019; Huang and Paul 2019). Their diversity arises embryonically from tightly regulated transcriptional programs which are spatially and temporally orchestrated in progenitor cells of the subpallium (Flames et al. 2007; Mayer et al. 2018; Mi et al. 2018). Progenitors from the medial ganglionic eminences (MGE) give rise to Parvalbumin (PV)-expressing basket and chandelier cells and to Somatostatin (SST)-expressing Martinotti and multipolar cells (Nigro, Hashikawa-Yamasaki, and Rudy 2018; Wichterle et al. 2001; Liodis et al. 2007; Wonders et al. 2008; Butt et al. 2008). This broad class of MGE-derived INs is under the control of several transcriptional regulators, including *Nkx2*.*1* (Butt et al. 2008) and *Lhx6* (Liodis et al. 2007), which have been used to genetically fate-map MGE-derived types. On the other hand, the caudal ganglionic eminence (CGE) (Vucurovic et al. 2010; Murthy et al. 2014; Lee et al. 2010) and to a lesser extent the preoptic area (POA) (Gelman et al. 2009; Niquille et al. 2018) contribute to a second group of INs whose common feature is to express the serotonin receptor 3a (HTR3A) (Vucurovic et al. 2010; Murthy et al. 2014; Lee et al. 2010). Functional integration of INs in the cortical circuits depends on their proper laminar positioning and their ability to connect to appropriate local cellular partners (Fishell and Kepecs 2019; Lim, Mi, et al. 2018). To reach the cortex, INs first leave their subpallial progenitor domains (Hernandez-Miranda et al. 2011; van den Berghe et al. 2013; Flames et al. 2004; Nobrega-Pereira et al. 2008; Marin et al. 2001; Rakic et al. 2015) and migrate tangentially within migratory streams located in the subventricular zone (SVZ) and marginal zone (MZ) (Lavdas et al. 1999; Lim, Pakan, et al. 2018; Marin and Rubenstein 2001; Martini et al. 2009; Tanaka et al. 2009; Tiveron et al. 2006). They then exit tangential streams to distribute in the nascent cortex, where they are driven towards specific layers and connect with preferential targets (Bartolini et al. 2017; Miyoshi and Fishell 2011; Sanchez-Alcaniz et al. 2011; Lopez-Bendito et al. 2008). Spatiotemporal patterning of subpallial progenitor domains plays an important role in determining the final laminar allocation of cortical INs. Indeed, MGE-derived INs mostly distribute in deep cortical layers (Pla et al. 2006), whereas HTR3A^+^ INs (hINs) originating from the POA/CGE mainly populate superficial layers (Vucurovic et al. 2010; Miyoshi et al. 2010; Lee et al. 2010; Murthy et al. 2014). In addition to these cell-intrinsic mechanisms, glutamatergic projection neurons (PNs) are able to influence the lamination of cortical INs. Namely, ectopic deep layer PNs can recruit LHX6^+^ INs (Lodato et al. 2011), while *Satb2*-expressing interhemispheric PNs from the visual cortex control cortical integration of hINs (Wester et al. 2019). How PNs settling into the cortical plate attract migrating INs has been well documented for MGE-derived INs (Bartolini et al. 2017) but the molecular mechanisms that regulate their laminar allocation is yet poorly elucidated. In addition to the attractive force that PNs of the cortical plate may exert on hINs, we present here a complementary mechanism in which MGE-derived INs constrain hINs into superficial layers. Here we find that the guidance receptor PlexinA4 (PLXNA4) is upregulated in hINs during their last phase of migration into the neocortex. We demonstrate that PLXNA4 conveys repulsive signaling of Semaphorin3A (SEMA3A), a guidance cue secreted by MGE-derived INs settled in deep cortical layers during perinatal days. Targeted deletion of SEMA3A in MGE-derived cells weakens the predilection for hINs to superficial layers, which in turn, distribute more in deep layers.

## Results

### Plexina4 *is upregulated in hINs during cortical plate invasion*

In a previous study we showed that the HTR3A regulates cortical invasion and laminar allocation of hINs (Murthy et al. 2014). Here we aimed to identify genes downstream of *Htr3a* that might specifically control the lamination of hINs. To this end, we used the *Gad65*-GFP mouse line, previously described as preferentially labeling hINs (Lopez-Bendito et al. 2004; Murthy et al. 2014). GFP^+^ cells were isolated by FACS from *Htr3a*^*+/+*^;*Gad65*-GFP^+^ and *Htr3a*^*-/-*^;*Gad65*-GFP^+^ cortices at three time-points reflecting the sequential process of hINs migration: E14.5 during tangential migration, E18.5 when invasion of the cortical plate (CP) takes place and P2 for laminar allocation. At E14.5, the whole cortical thickness was dissected to capture cells tangentially migrating along the MZ and IZ/SVZ streams (Fig. 1A left). At E18.5 and P2, migratory streams and the WM were discarded from the microdissection, in order to avoid hINs still migrating tangentially and isolate only those invading the CP (Fig. 1A middle and right). Gene expression was determined by microarray and levels of genes which were specifically upregulated during CP invasion in *Htr3a*^*+/+*^ hINs were compared to those expressed in *Htr3a*^*-/-*^ hINs. Among the 14 top enriched genes previously found to be specifically upregulated during CP invasion in *Htr3a*^*+/+*^ hINs (Murthy et al. 2014), we identified 5 genes (i.e., *Htr3a, Nr4a3, Plxna4, Dlg2* and *Cacna2d*) which failed to upregulate at E18.5 in *Htr3a*^*-/-*^ hINs (Fig. 1B right). Expression of the gene coding for the receptor PlexinA4 (*Plxna4*) had a similar dynamic expression to *Htr3a* (Fig. 1B left) and was of a particular interest since it has been shown to regulate neural cell guidance (Matsuoka et al. 2012; Hatanaka et al. 2019; Waimey et al. 2008; Okada and Tomooka 2012, 2013; Paap et al. 2016). Hypothesizing a role for *Plxna4* in the guidance and lamination of hINs, we first verified its expression in hINs along their migration to the cortex and as they transited through the different cortical compartments towards their final laminar positions. To do this, we performed fluorescent *in situ* hybridization (ISH) and analyzed levels of *Plxna4* expression in GFP^+^ cells from the *Htr3a*-GFP mouse line. At E14.5, *Plxna4* transcripts were almost undetectable in hINs tangentially migrating along the IZ/SVZ and MZ streams (Supp. Fig. 1A left). At E18.5, *Plxna4* expression was gradually increased in hINs as they left the tangential streams and migrated into the CP (Fig. 1C). A similar pattern was found at P2 when hINs were undergoing their terminal translaminar movements (Supp. Fig. 1A right). These data indicate that *Plxna4* is a *Htr3a*-dependent gene upregulated in hINs as they leave tangential streams to migrate into the CP.

**Figure 1.**
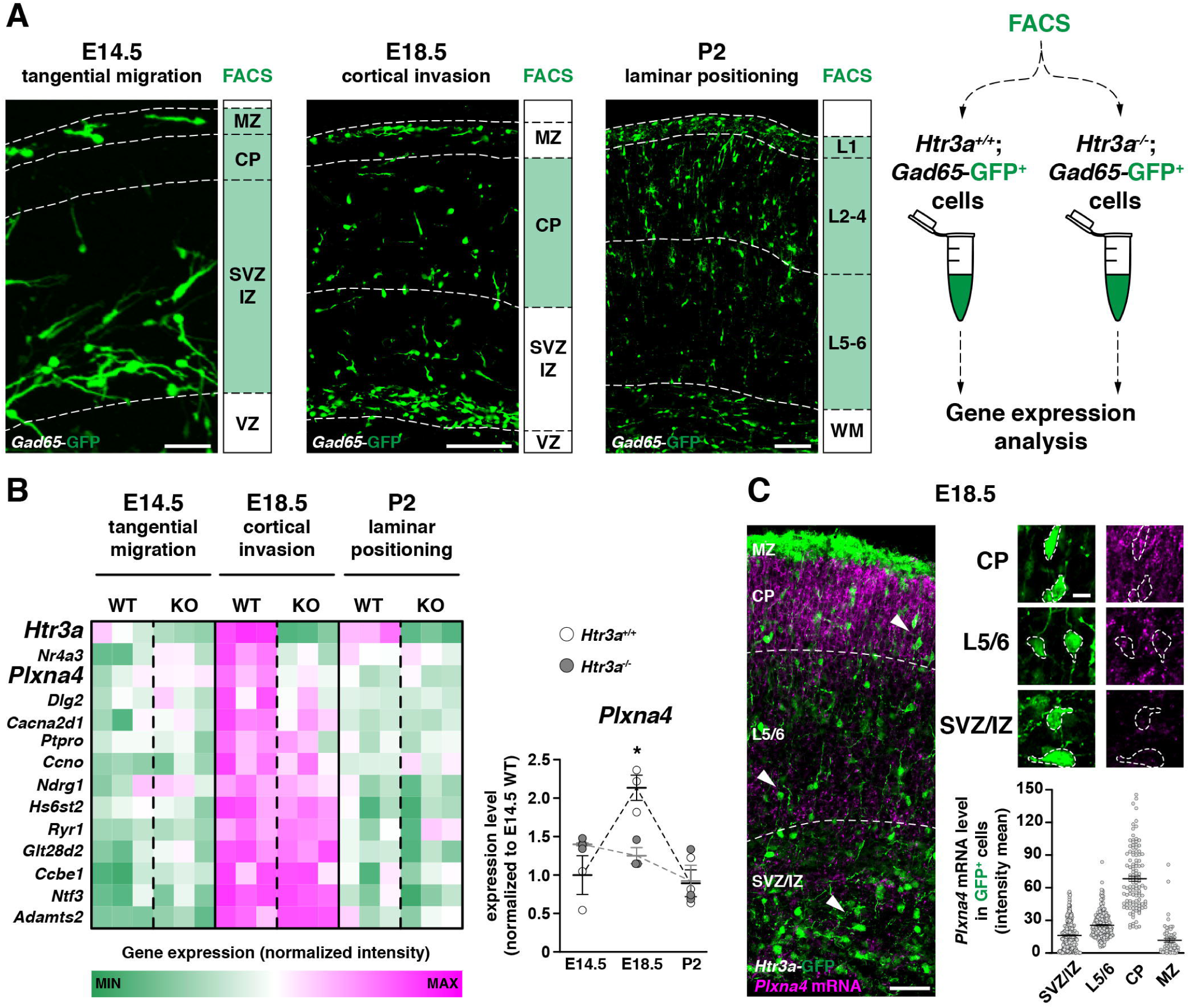
HTR3A^+^ interneurons (hINs) upregulate *Plxna4* during cortical plate (CP) invasion in a *Htr3a*-dependent manner. (A) Gene expression analysis using microarrays was performed on *Htr3a*^*+/+*^;*Gad65*-GFP^+^ and *Htr3a*^-/-^;*Gad65*-GFP^+^ cortical interneurons (INs) isolated by fluorescence assisted cell sorting (FACS) after microdisssection at E14.5, E18.5 and P2. White zones were discarded from the microdissection. Scale bar: 50µm (left), 100µm (middle and right) (B) Heat map showing top genes upregulated during CP invasion in *Htr3a*^+/+^;*Gad65*-GFP^+^ INs (WT) and expression of the same genes in *Htr3a*^-/-^;*Gad65*-GFP^+^ INs (KO). *Htr3a, Nr4a3, Plxna4, Dlg2* and *Cacna2d1* are not upregulated at E18.5 in *Htr3a*^-/-^;*Gad65*-GFP^+^ INs. *Plxna4* is specifically upregulated during CP invasion (E14.5 *vs* E18.5: **p = 0.0092), but this does not occur in the *Htr3a*^-/-^ condition (E14.5 *vs* E18.5: p = 0.5713; E18.5: *Htr3a*^+/+^ *vs Htr3a*^-/-^: fold change = 1.8; *p = 0.0415; Kruskal-Wallis and Dunn’s post-tests; n = 3 experiments for each condition). (C) *In situ* hybridization showing that at E18.5 *Plxna4* is expressed in hINs (white cell contours) having reached the CP. Fluorescence intensity cell mean indicates gradually increasing expression of *Plxna4* in hINs from tangential streams towards the CP at E18.5 (n = 2 brains, 341 cells in SVZ/IZ, 264 cells in L5/6, 117 cells in CP, 62 cells in MZ). Arrowheads point at high magnification positions. Scale bar: 50 µm (low magnification), 10 µm (high magnifications). Data are mean ± SEM. MZ: marginal zone, CP: cortical plate, SP: subplate, SVZ/IZ: subventricular/intermediate zone, VZ: ventricular zone, WM: white matter.

### Plxna4 *regulates the laminar allocation of hINs in the cortex*

To determine whether *Plxna4* regulates the laminar positioning of hINs *in vivo*, we analyzed their distribution in the somatosensory cortex of *Plxna4*^*-/-*^ mice and their littermate controls. We first aimed at ruling out potential cytoarchitectural defects in *Plxna4*^*-/-*^ brains. Although *Plxna4* has been involved in radial migration of PNs in rodents (Chen et al. 2008; Paap et al. 2016; Hatanaka et al. 2019), at P21 the overall cortical architecture and distribution of superficial layer PNs in *Plxna4*^-/-^ brains appeared unchanged as compared to controls (Supp. Fig. 2A). Secondly, although deletion of *Plxna1* has been shown to alter the proliferation of neuronal progenitors (Andrews et al. 2016), we observed here that the overall density of hINs was comparable in the two conditions at P21 (Fig. 2A, top), suggesting that neither the production nor the survival of hINs was affected by *Plxna4* deletion. Consistent with this observation, and with the barely detectable level of *Plxna4* expression found in tangentially migrating hINs, no differences were measured in the density of hINs in the migratory streams of *Plxna4*^*-/-*^ brains when compared to controls (Supp Fig. 2B). We next assessed the laminar allocation of hINs in *Plxna4*^*-/-*^ brains at P21 and found that they were shifted towards superficial cortical layers (L1-3) at the expense of L4 and deep layers (L5/6) (Fig. 2A, bottom). In contrast, the distribution of MGE-derived PV^+^ and SST^+^ INs remained unchanged (Fig. 2B, C). As the transcription factors PROX1, NR2F2 and SP8 were shown to contribute to the migration and development of CGE-derived cortical INs (Tang et al. 2012; Wei et al. 2019; Ma et al. 2012; Miyoshi et al. 2015; Kanatani et al. 2008), we examined whether the misplacement of hINs in *Plxna4*^*-/-*^ brains was associated to a change in their molecular identity. We focused on PROX1 because no clear changes have been reported in the laminar distribution of hINs subtypes following targeted deletion of SP8 (Wei et al. 2019; Ma et al. 2012) or NR2F2 (Tang et al. 2012) in INs. Using IHC, we found that the fraction of hINs expressing PROX1 was comparable between conditions (Supp. Fig. 2C), suggesting that deletion of *Plxna4* did not modify the molecular identity of hINs. Overall, these results indicate that *Plxna4* deletion does not affect the early steps of hINs tangential migration nor the laminar positioning of MGE-derived INs and PNs, but selectively alters the final laminar allocation of hINs, without altering their transcriptional features.

**Figure 2.**
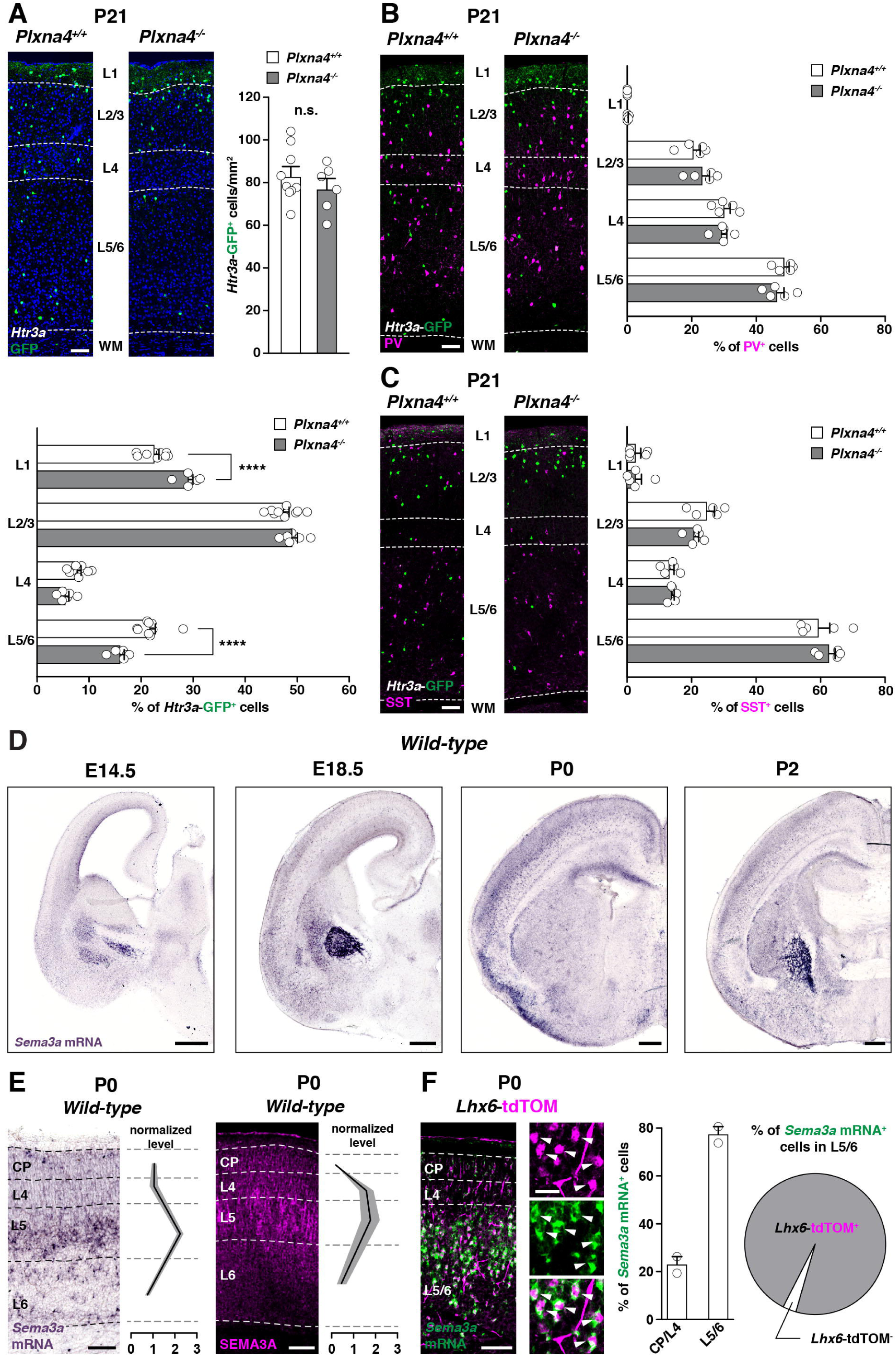
*Htr3a*-GFP^+^ interneurons (hINs) are mispositioned in *Plxna4*^*-/-*^ brains and MGE-derived interneurons (INs) are the perinatal cortical source of Semaphorin3A (SEMA3A) (A) Coronal slices of *Plxna4*^*+/+*^;*Htr3a*-GFP and *Plxna4*^*-/-*^;*Htr3a*-GFP brains showing positioning of hINs at P21 in the somatosensory cortex. The overall density of hINs (top) is similar in *Plxna4*^*+/+*^ and *Plxna4*^*-/-*^ cortices (*Plxna4*^*+/+*^: n = 9 brains, 4445 cells; *Plxna4*^*-/-*^: n = 6 brains; 3588 cells; p = 0.5287; Mann-Whitney test). The laminar positioning of hINs (bottom) is significantly altered in *Plxna4*^*-/-*^ brains when compared with *Plxna4*^*+/+*^ (*Plxna4*^*+/+*^: n = 9 brains, 8166 cells; *Plxna4*^*-/-*^: n = 6 brains, 6072 cells; ****p < 0.0001; 2-way ANOVA with Bonferroni’s post-tests). Scale bar: 100µm. (B-C) Immunohistochemistry (IHC) against the MGE-derived IN markers parvalbumin (PV) and somatostatin (SST) on coronal slices of *Plxna4*^*+/+*^ and *Plxna4*^*-/-*^;*Htr3a*-GFP brains at P21 in the somatosensory cortex. The distribution of PV^+^ (B, right) and SST^+^ (C, right) INs is not affected in *Plxna4*^*-/-*^ brains (PV: n = 5 brains, 2849 cells in *Plxna4*^*+/+*^, 2697 cells in *Plxna4*^*-/-*^; n.s.: p = 0.363; SST: n = 5 brains, 1825 cells in *Plxna4*^*+/+*^, 1563 cells in *Plxna4*^*-/-*^; n.s.: p = 0.2877; 2-way ANOVA). Scale bar: 100µm. (D) *In situ* hybridizations (ISH) for *Sema3a* on coronal sections at E14.5, E18.5, P0 and P2. Scale bar: 200µm. (E) *Sema3a* transcript and protein highlighted respectively by ISH (left) and IHC (right) on coronal sections at P0. Graphs of normalized intensity show that *Sema3a* is mainly expressed in L5 (left) from which the protein (right) distribute in gradient (ISH: n = 3 brains; IHC: n = 3 brains). Scale bars: 100µm. (F) ISH on coronal section of *Lhx6-*tdTOM brains at P0 showing that *Sema3a* mRNA is expressed in cells located in deep cortical layers 5/6 (middle; L5/6: 77.18 ± 3.43 %) and that these cells are MGE-derived INs (right; 96.94 ± 0.69 %; n = 2 brains, 730 cells). Scale bar: 100µm. All data are mean ± SEM. CP: cortical plate, WM: white matter.

### MGE-derived INs are the source of Semaphorin3A (SEMA3A) in deep layers

We next searched for ligands which could interact with PLXNA4. It has been previously shown that PLXNA4 conveys SEMA3A repulsive force during several processes of neuronal development (Battistini and Tamagnone 2016; Masuda and Taniguchi 2016; Pasterkamp 2012). Taking advantage of the Allen Brain Atlas for Developmental Brain database, we observed that the expression onset of *Sema3a* in the developing cortex occurred in the same temporal window as *Plxna4* upregulation in hINs, making it a suitable potential signaling partner. ISH for *Sema3a* performed at several developmental time-points (i.e., E14.5, E18.5, P0 and P2) confirmed its appearance in the cortex as of E18.5 and revealed that *Sema3a* expression was at its highest level at P0 (Fig. 2D). In previous RNA-sequencing analysis on INs at E18.5, *Sema3a* was strongly enriched (over 5-fold) in MGE-derived cells as compared to CGE-derived ones (Miyoshi et al. 2015). In agreement with this, we observed that, at P0, *Sema3a* mRNA was expressed in cells located in deep cortical layers (L5/6) (Fig. 2E, left), where MGE-derived INs mainly settle. The enrichment of SEMA3A was also validated at the protein level, with a highest expression level in L5 (Fig. 2E, right). To determine whether MGE-derived INs are the source of SEMA3A, we crossed *Lhx6*^cre^ driver mice with *Rosa26*-tdTOM^*fl/fl*^ reporters (*Lhx6*-tdTOM) to map MGE-derived interneurons and verified the specificity of recombination (Supp. Fig. 2D, E). Analysis of brain sections from *Lhx6*-tdTOM mice (Fig 2F) confirmed that the majority of *Sema3a*-expressing cells were located in deep layers (∼80%) and that virtually all of them were corresponding to MGE-derived INs (>95%). A similar analysis was performed on *Htr3a*-GFP brain sections and showed that only a minimal fraction (∼10%) of hINs was expressing *Sema3a* (Supp. Fig. 2F). These results show that MGE-derived INs settling in deep layers generate a local source of SEMA3A in the forming cortex.

### SEMA3A induces growth cone collapse of hINs through a PLXNA4-NRP1 mechanism

We then addressed the question of a potential SEMA3A interaction with *Plxna4*-expressing hINs. As a prerequisite for SEMA3A signaling through PLXNA4, we first verified that neuropilin (NRPs) coreceptors were expressed by hINs (Fig. 3A top). To this purpose, we performed ICC on E14.5 dissociated cells from the CGE cultured for 3 days *in vitro* (3DIV) (Fig. 3A bottom) and found that hINs expressed PLXNA4 along with NRP1 and NRP2, and that these proteins colocalized on the cell body and growth cone (GC) (Fig. 3B). To evaluate GC response of hINs to SEMA3A, cell cultures from *Plxna4*^*+/+*^;*Htr3a*-GFP and *Plxna4*^*-/-*^;*Htr3a*-GFP CGEs were challenged with either 5nM SEMA3A-Fc chimera or vehicle solutions (Fig 3C). SEMA3A application significantly decreased the size of GCs in *Plxna4*^*+/+*^;*Htr3a*-GFP^+^ INs as compared to the vehicle condition (Fig. 3C, Supp. Fig. 3A). Notably, this effect was absent in *Plxna4*^*-/-*^*;Htr3a*-GFP^+^ INs, indicating that PLXNA4 is required for SEMA3A to induce a GC collapse (Fig 3C, Supp. Fig. 3A). Furthermore, by preventing SEMA3A binding using blocking peptides (Fig. 3A top, D), we found that NRP1 but not NRP2 inactivation rescued the SEMA3A-induced decrease in GC size of *Htr3a*-GFP^+^ INs (Fig 3D, Supp. Fig. 3B). Taken together, these *in vitro* experiments demonstrate that extracellular SEMA3A elicits GC collapse of hINs via the PLXNA4/NRP1 receptor complex.

**Figure 3.**
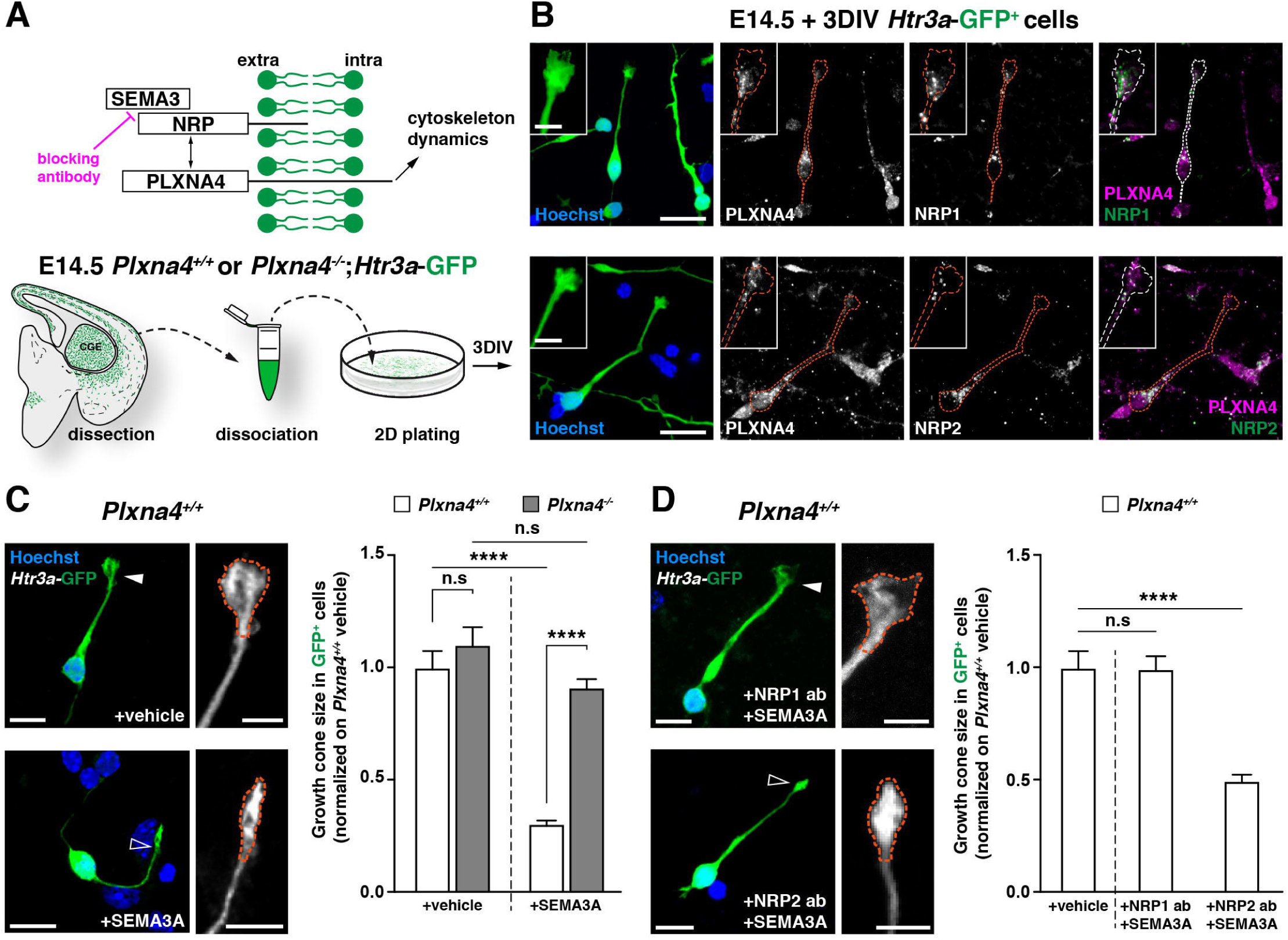
SEMA3A induces growth cone collapse of *Htr3a*-GFP^+^ interneurons (hINs) through the PLXNA4/NRP1 complex. (A) Schematic representation (top) showing that extracellular class 3 SEMA binds to Neuropilin (NRP) receptor, which forms a complex with PlexinA4 (PLXNA4). Upon binding, intracellular domain of PLXNA4 unfolds to activate a downstream pathway for cytoskeletal reorganization. Paradigm for hINs dissociated cell culture preparations (bottom). (B) Immunocytochemistry on E14.5 cells dissociated from the CGE and cultured for 3 days *in vitro* (3DIV) showing that hINs coexpress neuropilin 1/2 (NRP1/2) and PLXNA4 both in the soma and growth cones (GCs, insets). Scale bar: 20µm, 10µm (insets). (C) Developed (top, arrowheads) and collapsed (bottom, open arrowheads) GCs of *Plxna4*^*+/+*^ hINs after vehicle and SEMA3A application, respectively. GC size of *Plxna4*^*+/+*^ and *Plxna4*^*-/-*^ hINs after vehicle application were comparable (*Plxna4*^*+/+*^: n = 3 experiments, 63 GCs; *Plxna4*^*-/-*^: n = 5 experiments, 105 GCs; n.s.: p > 0.99). SEMA3A application induces GC collapse of *Plxna4*^*+/+*^ hINs (vehicle: n = 3 experiments, 63 GCs; SEMA3A: n = 7 experiments, 427 GCs; ****p < 0.0001) but not of *Plxna4*^*-/-*^ hINs (n = 5 experiments, 365 GCs; n.s.: p = 0.8055). Statistics are done with Kruskal-Wallis test with Dunn’s post-tests. Scale bar: 20µm, 10µm (insets). (D) Examples of GC after SEMA3A application with NRP1 (top) or NRP2 (bottom) blocking peptides. Collapse rescue is achieved by blocking NRP1 but not NRP2. (NRP1: n = 4 experiments, 176 GCs; NRP2: n = 4 experiments, 244 GCs; n.s.: p > 0.99 and ****p < 0.0001, respectively; Kruskal-Wallis test with Dunn’s post-test). Scale bar: 20µm, 10µm (insets). Data are mean ± SEM normalized with *Plxna4*^*+/+*^ GC size mean in vehicle condition.

### SEMA3A is chemorepulsive for migrating hINs

We next investigated the outcome of SEMA3A/PLXNA4 signaling in a model closer to the *in vivo* situation and therefore tested whether SEMA3A induced GC collapse of hINs *ex vivo*. To this purpose, E17.5 acute coronal slices were prepared from *Plxna4*^*+/+*^;*Htr3a*-GFP and *Plxna4*^*-/-*^;*Htr3a*-GFP brains and GCs of GFP^+^ INs migrating in the CP were imaged along a time period of 10min vehicle or 100nM SEMA3A application, followed by 72min ASCF superfusion (Fig. 4A top). At the beginning of the imaging, the average intensity (here used as proxy for the filopodia/lamellipodia state) (Fig. 4A bottom) and the size of the GC were comparable in all experimental conditions (Supp. Fig. 4A). However, as considerable variability was observed in GC size, its variation along the imaging session was measured as a percentage of the starting size to capture at best the effect of the SEMA3A on individual GC (Fig. 4B left, Supp. Fig. 4B left). While the GC area remained overall constant with vehicle application, SEMA3A repeatedly induced a reduction in GC size of *Plxna4*^*+/+*^;*Htr3a*-GFP^+^ INs (Fig 4B left). By contrast, SEMA3A superfusion did not induce size changes in GCs of *Plxna4*^*-/-*^;*Htr3a*-GFP^+^ INs (Fig 4B left). Paired measurements of the GC size and average intensity in initial (P1) and final (P2) periods showed that both parameters were significantly changed upon SEMA3A application in *Plxna4*^*+/+*^;*Htr3a*-GFP^+^ INs but not in *Plxna4*^*-/-*^;*Htr3a*-GFP^+^ INs (Fig 4B right, Supp. Fig. 4B right). These results indicate that SEMA3A is able to collapse GC of hINs *ex vivo* and that this effect is PLXNA4 dependent. Because GC collapse is often associated with a subsequent avoidance reaction to a cue, we investigated whether SEMA3A could act as a chemorepellent for hINs as they invade the nascent cortex. To test this hypothesis, we assessed the migratory behavior of hINs as they encountered a source of SEMA3A. Specifically, we transfected HEK-T293 cells with either *pU6*-tdTOM (control) or *pU6*-*Sema3a*-tdTOM expression constructs and engrafted these cells into the CP of *Htr3a*-GFP^+^ acute slices at E17.5 (Fig 4C right). Migratory movements of *Htr3a*-GFP^+^ INs in the vicinity of engrafted HEK-T293 cells were assessed during a 10h-long live imaging session (Fig 4C left). Analyses indicated that the density of hINs increased similarly around the control and SEMA3A-expressing explants during the first 3h. Strikingly, at later time-points, hINs density in the 50µm area surrounding the explant was significantly lower in the SEMA3A overexpression condition as compared to control (Fig 4D left, Supp. Fig. 4C). This difference was maintained along a plateau of 7h, precluding a delayed invasion around the SEMA3A-expressing explant. Although statistically non-significant, a similar effect was observed in a more distant area (50-100µm) surrounding the explant, suggesting that SEMA3A signaling was dose-dependent (Fig 4D left, Supp. Fig. 4C, Supp. movie 2). Finally, we observed that in cortices engrafted with control tdTOM-expressing HEK-T293 cells, hINs migrated into the explant, while the number of these events was significantly reduced in the SEMA3A overexpression condition (Fig 4D right). Overall, these results indicate that SEMA3A acts as chemorepellent for migrating hINs, most probably through PLXNA4-dependent collapse of their GCs.

**Figure 4.**
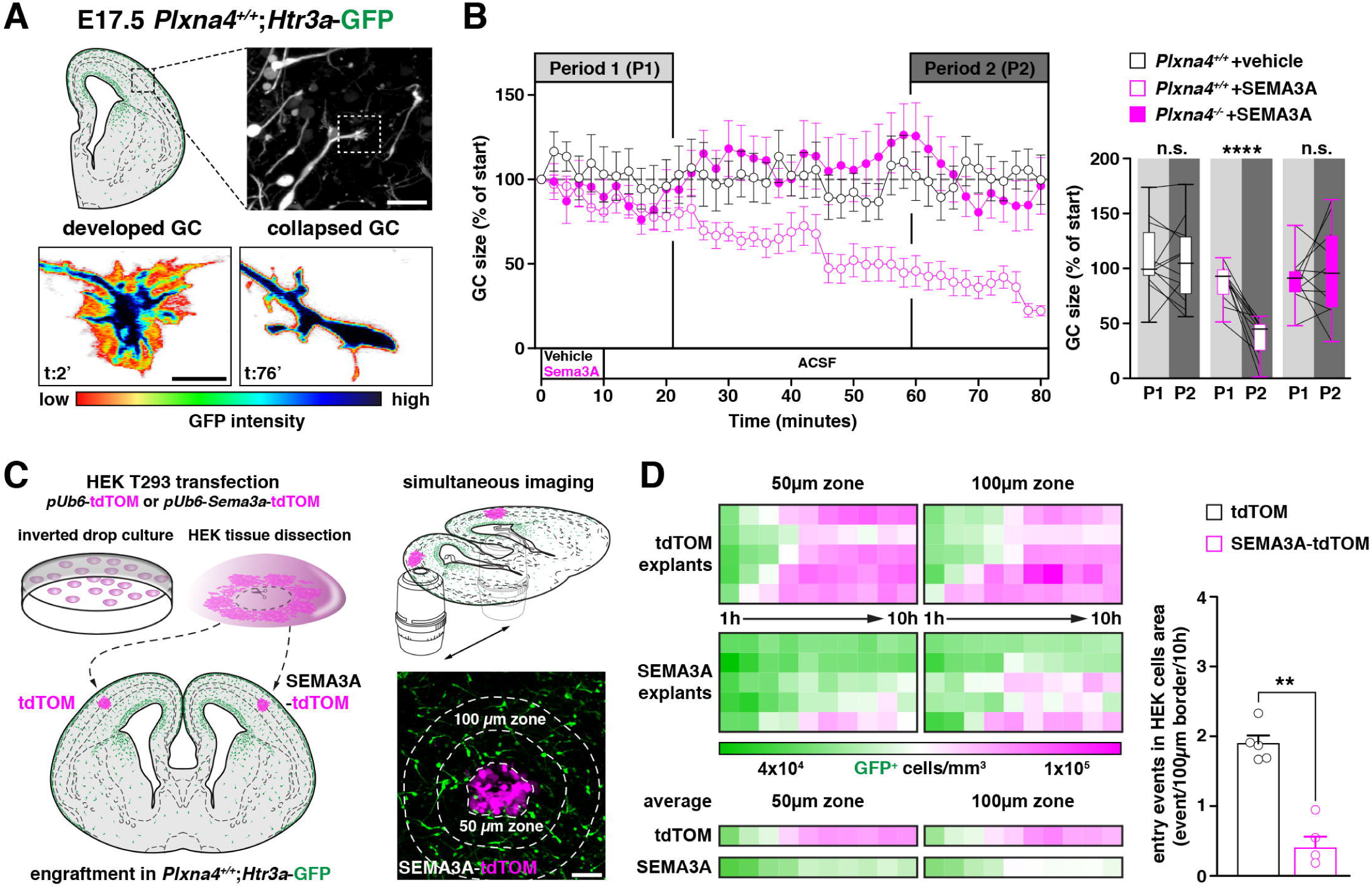
SEMA3A is chemorepulsive for migrating *Htr3a*-GFP^+^ interneurons (hINs) (A) Schematic of *Htr3a*-GFP^+^ coronal section at E17.5 indicating that live imaging is focused on GCs of hINs migrating in the CP (top). Images (bottom) showing the developed (left) and collapsed (right) morphology of a single GC. The endogenous GFP intensity (pseudocolor) is used as a proxy to assess the state of GC structures. Scale bar: 50µm (top), 10µm (bottom). (B) GC size is measured every 2 min along a timeline consisting of 10 min vehicle (black) or SEMA3A (magenta) application, followed by 72 min of ACSF superfusion (left). Graph of GC size (percentage of the starting value) shows that GCs of *Plxna4*^*+/+*^ hINs (n = 12; black open circles) are relatively constant in size along vehicle perfusion, but collapse within an hour following SEMA3A application (n = 11; magenta open circles). GCs of *Plxna4*^*-/-*^ hINs are not affected by SEMA3A (n = 11; magenta circles). Graphs of paired values (right) from the initial period (P1; 0-20 min) and the final period (P2; 60-80 min) shows that vehicle does not impact GCs of *Plxna4*^*+/+*^ hINs (n.s.: p = 0.8728; paired t-test) whereas SEMA3A dramatically reduced their size (****p < 0.0001; paired t-test). GCs of *Plxna4*^*-/-*^ hINs are not affected by SEMA3A (n.s.: p = 0.5504; paired t-test). (C) Paradigm of SEMA3A ectopic expression in *ex vivo* experiments. HEK-T293 pseudotissue transfected with either *pUb6*-tdTOM (i.e., control) or *pUb6-SEMA3A*-tdTOM plasmids is manually dissected and transplanted in the cortical area of single *ex vivo* slice from E17.5 *Plxna4*^*+/+*^;*Htr3a*-GFP brains (left). Grafted areas are simultaneously image every 10min over 10h (top right) and hINs density is measured every hour in a 50µm and 100µm zone around the HEK-T293 pseudotissue (bottom right). (D) Heatmaps (left) representing hINs density in the 50µm and 100µm zone for each slices (up) and their average (bottom). After 4h, a significantly smaller hINs density is observed in the 50µm area but not in the 100µm area around the SEMA3A-expressing explants compared to the control explants indicating a repulsive action of SEMA3A on *Plxna4*^*+/+*^ hINs (n = 5 explants for each condition, control explant: 9244 cells; SEMA3A explant: 5834 cells; 50µm area: *p < 0.01, **p < 0.001; 100µm area: n.s.: p > 0.05; 2-way ANOVA with Geisser Greenhouse correction and Tukey’s post-tests, see Supp. Fig. 4C). *Plxna4*^*+/+*^ hINs movements across the HEK-T293 border (right) occur more often in control (n = 81 events) compared to SEMA3A explants (n = 23 events) (**p = 0.0079; Mann-Whitney test). Scale bar: 50µm. Data in B (left) and D are mean ± SEM; whisker plots in B (right) represent 95% percentile with mean ± SD.

### SEMA3A-PLXNA4 interactions regulates the laminar allocation of hINs

We then wanted to further verify the repulsive action of SEMA3A on hINs *in vivo*. For this purpose, we induced an ectopic overexpression of SEMA3A in superficial layers (L2/3) where the majority of hINs normally settle. To label and manipulate PNs in L2/3, we performed *in utero* electroporation (IUE) targeting the dorsal wall at E15.0 (Fig. 5A). Efficient overexpression of *Sema3a* was validated in PNs of prospective L2/3 at E18.5 (Fig. 5B), when hINs were invading the CP. Overexpression of SEMA3A in L2/3 did not affect laminar distribution nor modify the position of electroporated PNs at P21 (Supp. Fig. 5A). By contrast, the fraction of hINs located in L2/3 significantly decreased as compared to the control condition. This was compensated by a significant increase of hINs located in L1 (Fig. 5C). As a consequence, the density of hINs located in the L2/3 electroporated region was significantly decreased in the SEMA3A condition as compared to control brains (Fig. 5E). The fact that hINs were shifted from L2/3 to L1, but not towards L5/6, suggested the additional repulsive action of SEMA3A endogenously found in deep cortical layers. To assess the requirement of PLXNA4 in this process, we replicated IUE of SEMA3A in *Plxna4*^-/-^;*Htr3a*-GFP embryos. In contrast to the *Plxna4*^*+/+*^ condition, the laminar distribution of hINs and their density in the L2/3 of *Plxna4*^-/-^ brains were not altered with overexpression of SEMA3A as compared to tdTOM (Fig. 5D, E), thus indicating that PLXNA4 was required for SEMA3A to repulse hINs. Altogether, these results reveal that SEMA3A exerts a repulsive action on hINs through a PLXNA4-dependent mechanism.

**Figure 5.**
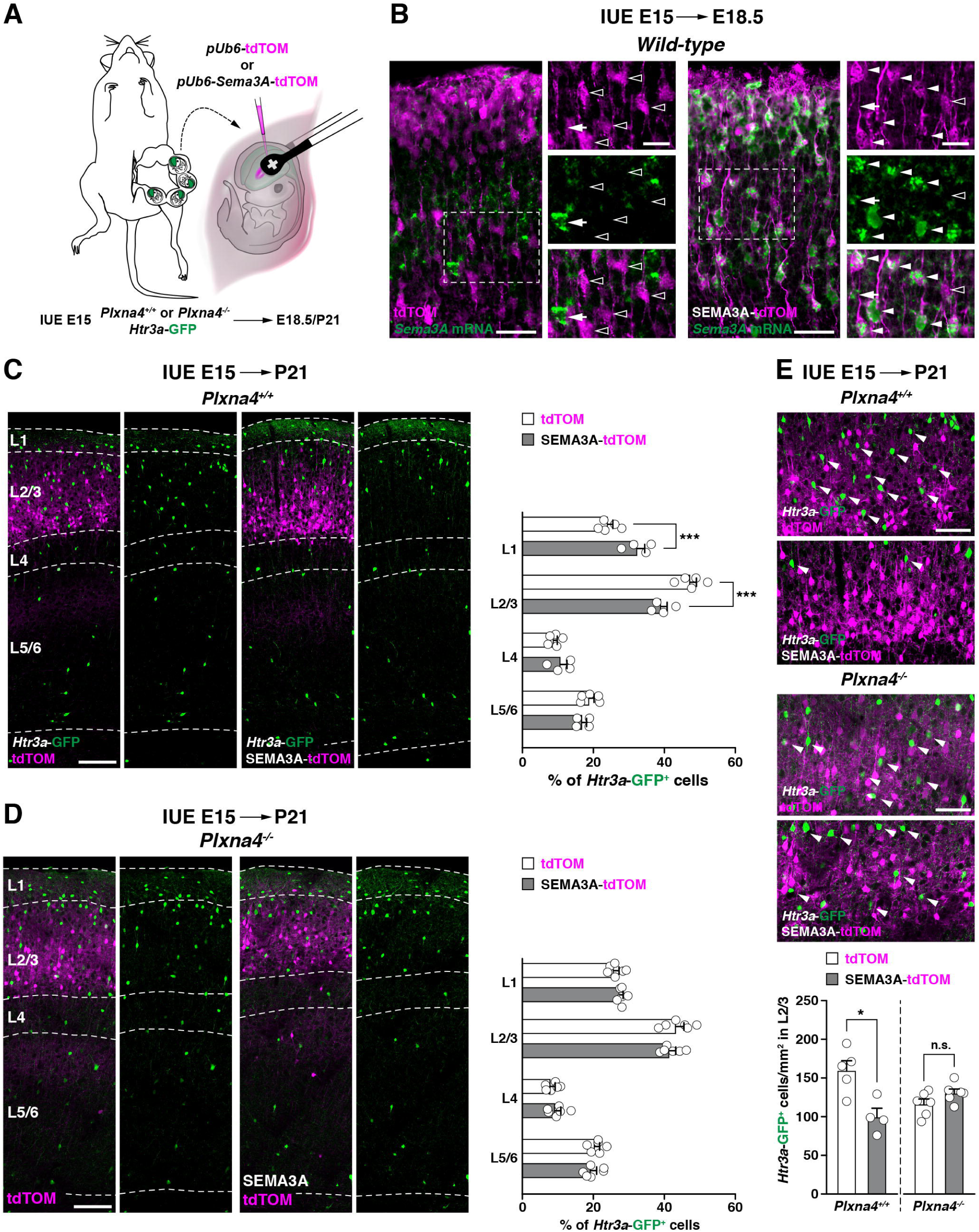
SEMA3A regulates the cortical laminar allocation of *Htr3a*-GFP^+^ interneurons (hINs) through PLXNA4. (A) Experimental paradigm of *in utero* electroporation (IUE). To manipulate pyramidal cells of prospective L2/3, the pallial wall of *Plxna4*^+/+^;*Htr3a*-GFP or *Plxna4*^-/-^;*Htr3a*-GFP embryos is electroporated at E15.0 with either *pUb6*-tdTOM (control) or *pUb6*-*Sema3a-*tdTOM plasmid. (B) Images showing E18.5 cortices from electroporated brains at E15.0. *In situ* hybridization confirms the overexpression of *Sema3a* in pyramidal neurons electroporated with *pUb6*-*Sema3a-*tdTOM (right, arrowheads), but not with in control (left, open arrowheads). Arrows point at endogenous *Sema3a*-expressing cells. Scale bars: 100µm (high magnification), 20µm (low magnification). (C-D) Illustrative images showing the distribution of *Plxna4*^+/+^ (C) or *Plxna4*^-/-^ hINs (D) in the P21 somatosensory cortices of brains electroporated at E15 with control or SEMA3A overexpressing construct. Layering analysis showing misdistribution of *Plxna4*^+/+^ hINs (C, left), but not *Plxna4*^-/-^ hINs (D, left) in upper cortical layers following SEMA3A overexpression (*Plxna4*^+/+^ control: n = 5 brains, 3425 cells; *Plxna4*^+/+^ SEMA3A: n = 4 brains, 1960 cells; L1: ***p = 0.0006; L2/3: ***p = 0.0007, 2-way ANOVA with Bonferroni’s post-test; *Plxna4*^-/-^ control: n = 6 brains, 3947 cells; *Plxna4*^-/-^ SEMA3A: n = 6 brains, 3508 cells; n.s.: p = 0.2325, 2-way ANOVA). Scale bar: 200µm. (E) High magnification on L2/3 IUE zone of *Plxna4*^+/+^ (top) or *Plxna4*^-/-^ (middle) brains after IUE with control or SEMA3A overexpressing construct. Cell density (bottom) of *Plxna4*^+/+^ hINs but not of *Plxna4*^-/-^ hINs is decreased in L2/3 following SEMA3A overexpression (*Plxna4*^+/+^ control: n = 5 brains, 1353 cells; *Plxna4*^+/+^ SEMA3A: n = 4 brains, 538 cells; *Plxna4*^-/-^ control: n = 6 brains, 1445 cells; *Plxna4*^-/-^ SEMA3A: n = 6 brains, 1132 cells; *p = 0.0317, n.s.: p = 0.1797, Mann-Whitney test). Scale bar: 100µm. All data are mean ± SEM.

### *Deletion of* Sema3a *in MGE-derived INs specifically affects the laminar distribution of hINs*

We finally hypothesized that MGE-derived INs regulate the positioning of superficial cortical hINs through the secretion of SEMA3A. Given that MGE-derived INs settle in the cortex before hINs (Batista-Brito and Fishell 2009), they could act as guidepost cells driving hINs towards superficial layers. To determine the responsiveness of hINs when encountering deep layers, endogenously enriched in SEMA3A, we developed an *ex vivo* model where SVZ streams from *Plxna4*^*+/+*^ or *Plxna4*^*-/-*^ brains at birth were engrafted in the WM of wild-type isochronic hosts (Supp. Fig. 6A). After 12h-long live imaging, we analyzed the dynamics of hINs exiting the engraftment towards the CP/superficial layers. Although the average speed of hINs in the IZ was comparable between the two conditions, *Plxna4*^*+/+*^ hINs moved significantly slower when entering the SEMA3A-enriched zone (Supp. Fig. 6B), suggesting a change in migratory behavior of PLXNA4-expressing hINs in response to endogenous SEMA3A signaling. In addition to the decrease in speed, we observed that *Plxna4*^*+/+*^ hINs were spending significantly less time in the SEMA3A-enriched zone when compared to *Plxna4*^*-/-*^ hINs (Supp. Fig. 6C). This result suggested that, while migrating from the IZ to the CP, hINs progressively gain responsiveness to SEMA3A, which kept them away from deep layers. In order to finally determine the role of SEMA3A in hINs neocortical settlement, we induced a targeted deletion of *Sema3*a in MGE-derived INs by crossing *Lhx6*^cre^ mice with *Sema3a*^*fl/fl*^;*Htr3a*-GFP mice (*Lhx6*^*Sema3a-/-*^;*Htr3a*-GFP) (Fig. 6A). With this strategy we aimed at specifically impeding *Sema3a* expression in LHX6^+^ cells, without affecting their migration and settlement into the neocortex. Indeed, ISH for *Lhx6* showed comparable patterns in *Lhx6*^*Sema3a+/+*^and *Lhx6*^*Sema3a-/-*^ brains at P0 (Fig. 6B left, arrowheads). As expected, the deletion was specific to *Lhx6*-expressing cells (Fig 6B right, open arrowheads) as hybridization signal for *Sema3a* mRNA was still present in *Lhx6*^*Sema3a-/-*^ brain regions were *Lhx6* mRNA was hardly detectable (Fig 6B). These observations confirmed that our model was selectively preventing production of *Sema3a* in *Lhx6*-expressing INs, without interfering with their genesis and migration into the neocortex. In addition, analysis at P21 confirmed that *Sema3a* deletion did not affect their final laminar positioning nor density (Fig. 6C) and did not alter the overall cortical architecture of superficial layer PNs (Supp. Fig. 6D). In striking contrast, whereas their density was comparable in both conditions, hINs were misallocated throughout cortical layers in *Lhx6*^*Sema3a*-/-^ mice as compared to controls (Fig. 6D). Higher fractions of hINs in *Lhx6*^*Sema3a-/-*^ brains were found to be allocated in L4 and deep layers L5/6 at the expense of superficial ones, indicating their likelihood to invade and settle in the area where SEMA3A no more exerted its repulsive action. Taken together, our data support a model (Fig. 7) where SEMA3A secreted by deep layer MGE-derived INs cell-extrinsically regulates the laminar allocation of superficial layer hINs through a SEMA3A-PLXNA4 interaction.

**Figure 6.**
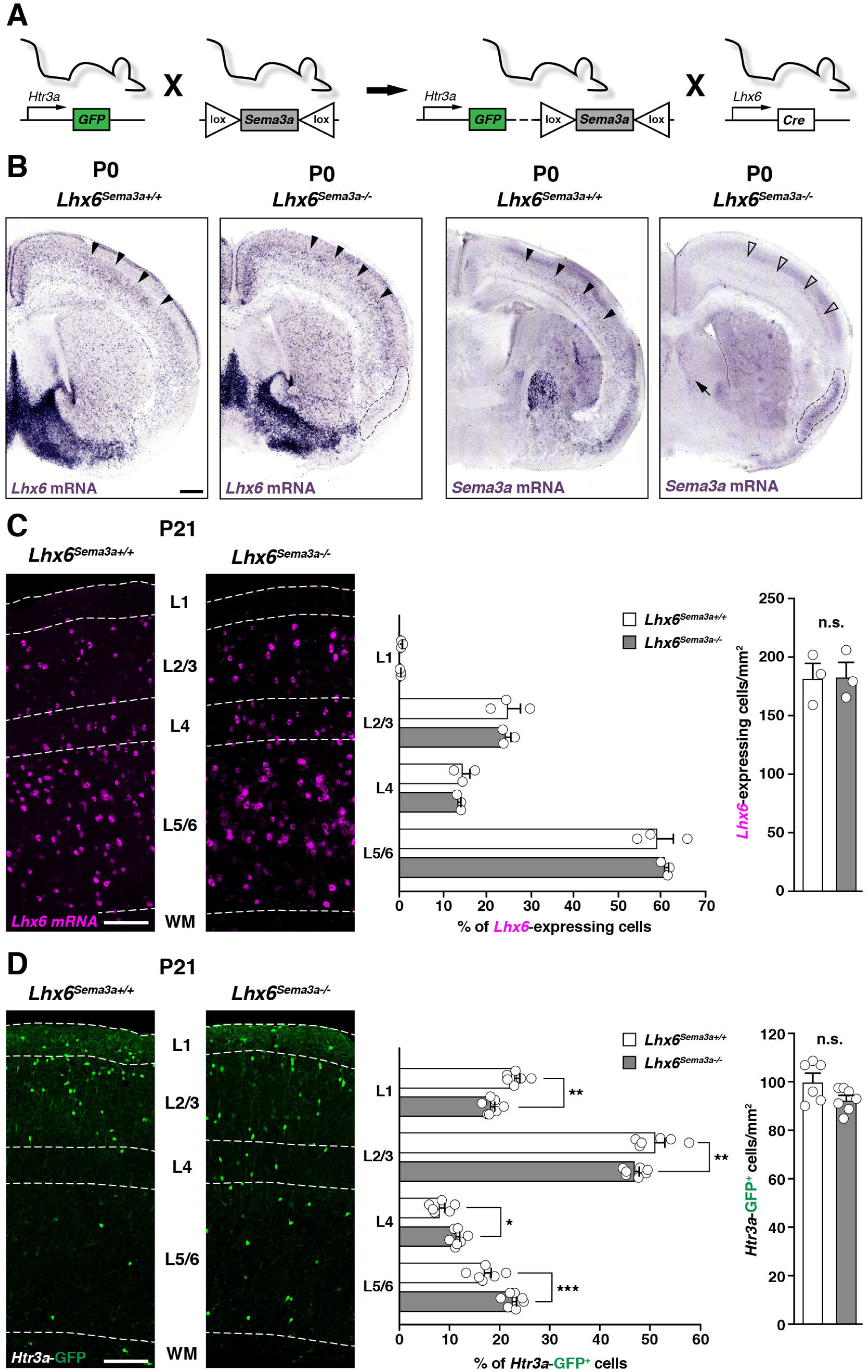
Deletion of *Sema3a* in MGE-derived interneurons (INs) alters the laminar positioning of *Htr3a*-GFP^+^ INs (hINs) (A) Paradigm of mice breeding to obtain *Sema3a* deletion in MGE-derived INs. (B) *In situ* hybridization for *Lhx6* (left) and *Sema3a* (right) in P0 coronal slice of *Lhx6*^*cre*^;*Sema3a*^+/+^ (*Lhx6*^*Sema3a+/+*^) and *Lhx6*^*cre*^;*Sema3a*^*fl/fl*^ mice (*Lhx6*^*Sema3*-/-^) mice. While *Lhx6* mRNA expression is unaffected, *Lhx6*^*Sema3*-/-^ mice show a total depletion of *Sema3a* mRNA in the neocortex (open arrowheads), but not in areas where its expression is not due to *Lhx6*-expressing cells (e.g., piriform cortex, dashed line and septum, arrowhead). Scale bar: 200µm. (C) Coronal slices showing the laminar distribution of *Lhx6*-expressing INs in *Lhx6*^*Sema3a*+/+^ and *Lhx6*^*Sema3*-/-^ somatosensory cortices at P21. The laminar positioning (middle) and density (right) of *Lhx6*-expressing INs is not modified in the two conditions (*Lhx6*^*Sema3a*+/+^: n = 3 brains, 4445 cells; *Lhx6*^*Sema3a*-/-^: n = 3 brains, 4754 cells; density: n.s.: p > 0.99; Mann-Whitney test; lamination: n.s.: p = 0.8959; 2-way ANOVA). Scale bar: 200µm (D) Coronal slices showing the laminar distribution of hINs in *Lhx6*^*Sema3a*+/+^ and *Lhx6*^*Sema3*-/-^ somatosensory cortices at P21. The laminar positioning of hINs (middle) is significantly altered in *Lhx6*^*Sema3*-/-^ cortices when compared with *Lhx6*^*Sema3a*+/+^ (*Lhx6*^*Sema3a*+/+^: n = 6 brains, 2829 cells; *Lhx6*^*Sema3a*-/-^: n = 7 brains, 6104 cells; L1: **p = 0.0016, L2/3: **p = 0.0074, L4: *p = 0.0397, L5/6: ***p = 0.0002; 2-way ANOVA with Bonferroni’s post-test). The overall density of hINs (right) is similar in *Lhx6*^*Sema3a*+/+^ and *Lhx6*^*Sema3*-/-^ somatosensory cortices (n.s.: p = 0.2343; Mann-Whitney test). Scale bar: 200µm All data are mean ± SEM. WM: white matter.

**Figure 7.**
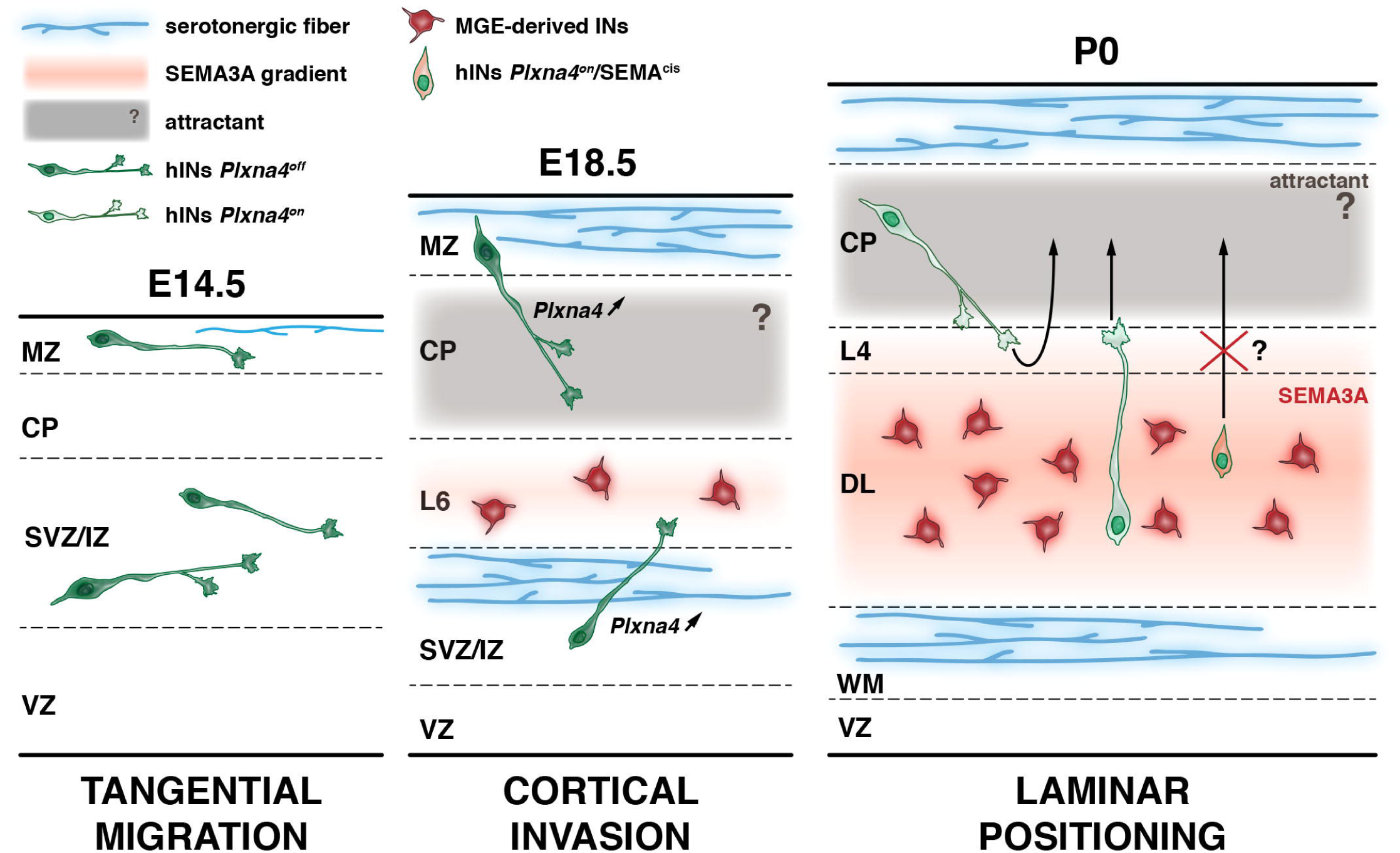
Model of hINs lamination by MGE-derived Ins. At E14.5, hINs are tangentially migrating in streams (dark green) while serotonergic afferent start invading the cortical area (blue). As hINs leave the streams and starts invading the cortical plate at E18.5, they upregulate *Plxna4*. This upregulation may be due to signals from the serotonergic system going through HTR3A receptors whose expression is concomitantly upregulated in hINs. At the same time, MGE-derived INs start expressing SEMA3A in deep layers of the forming cortex (red). The gradual increased expression of PLEXINA4 in hINs (light green) and SEMA3A in MGE-derived INs in deep layers (red) is repelling hINs towards superficial layer and precluding backward movements from hINs in superficial layers towards deep layers (P0). Although not discovered yet, an attractant produced by PNs of the CP may contribute to guide hINs into superficial layers (gray). The small population of hINs that settle in deep layers may be unsensitive to SEMA3A *trans* signaling from MGE-derived INs due to expression of *cis* SEMA3A (orange).

## Discussion

INs are essential to cortical function, managing the modulation of microcircuit activity. From their discrete sites of origin, INs engage in tangential migration to invade the forming neocortex, where hINs preferentially settle in superficial layers, while MGE-derived INs mainly populate deep ones. Here we identify the guidance receptor PLXNA4 as a mediator of hINs allocation into superficial layers. We find that *Plxna4* is downstream of *Htr3a*, and upregulated in hINs as they invade the CP. With gain and loss-of-function experiments, combined to *in vitro, ex vivo* and *in vivo* techniques, we show that migrating hINs are repelled by the secreted guidance cue SEMA3A and that this process is mediated by PLXNA4. Finally, we identify that MGE-derived INs, already settled in deep cortical layers, are the source of secreted SEMA3A and that targeted disruption of this cue affects hINs laminar distribution. Taken together, our results demonstrate that MGE-derived INs, the first in place, act as guidepost cells controlling the sorting of hINs into superficial layers. While PN-IN interactions have been the ground for recent advances, this work unveils an unknown and novel type of interaction in the field of interneuron guidance.

### SEMA3A as a cue regulating the allocation of cortical hINs

The diversity of cortical INs arises from early spatial and temporal patterning events (Fishell and Kepecs 2019; Lim, Mi, et al. 2018). Transcriptional programs governing their specification and differentiation predispose them to allocate in distinct cortical layers (Liodis et al. 2007; Miyoshi et al. 2015; Miyoshi and Fishell 2011; Mayer et al. 2018; Wamsley and Fishell 2017). Although cell-intrinsic transcriptomic programs are key determinant in these processes, the local environment provides a necessary modulation through context-dependent mechanisms (Fishell and Kepecs 2019). The cellular sources of these cell-extrinsic signals are likely to be diverse. Among the most studied, PNs located in deep and superficial cortical layers regulate the differential laminar distribution of cortical INs (Lim, Mi, et al. 2018; Lodato et al. 2011; Wester et al. 2019), but the molecular mechanisms through which distinct sets of PNs control the allocation of different types of cortical INs into specific cortical layers remains quite unexplored. It has been proposed that INs and PNs, eventually paired into cortical circuits, express complementary molecules which ultimately control the lamination of INs. Among potential cues secreted by PNs, Neuregulin 3 (NRG3) has been shown to act as a chemoattractant, thus facilitating the invasion of *Erbb4*-expressing MGE-derived cortical INs into the CP (Bartolini et al. 2017). Here we reveal a novel cellular source providing a guidance cue that influences the lamination of hINs during the perinatal period. In accordance with previous works (Mayer et al. 2018; Miyoshi et al. 2015), we find that SEMA3A is strongly enriched in MGE-derived INs while POA/CGE-derived hINs are still in the process of actively migrating towards the CP. We hypothesized that, given its localized expression in deep layers, SEMA3A instructs hINs to preferentially settle in superficial layers. Concordantly, with *ex vivo* experiments on acute slices we demonstrate that SEMA3A induces GC collapse and acts as a non-permissive/repulsive cue on hINs. Furthermore, we find that the generation of an ectopic source of SEMA3A in superficial layer PNs is able to persistently misplace hINs from their usual location in the cortex. Given these findings, we propose that an INs-INs interaction supplements PNs-INs signaling and participates in hINs lamination.

### SEMA3A-PLXNA4 interaction controls hINs lamination

The serotonergic system controls the migration of hINs into the CP and their lamination by acting through HTR3A (Frazer, Otomo, and Dayer 2015; Murthy et al. 2014). Here we show that in hINs *Plxna4* upregulation during the CP invasion is dependent of *Htr3a*, suggesting a link between the serotonergic system and further events of hINs guidance relying on PLXNA4. PLXNA4-related mechanisms have been extensively described for axon guidance (Mitsogiannis, Little, and Mitchell 2017; Runker et al. 2008; Schwarz et al. 2008; Suto et al. 2005; Yaron et al. 2005) as well as in glial (Okada and Tomooka 2012, 2013) and neuronal migration (Paap et al. 2016). In the field of axonal guidance, it has been proposed that SEMA3A has a favored affinity for NRP1 which in turn pairs principally with PLXNA4, whereas PLXNA3/NRP2 complexes transduce SEMA3F signaling (Suto et al. 2005; Yaron et al. 2005). This concept of preferential association between PLXNA4 and PLXNA3 with NRP1 and NRP2 respectively is also found in the field of neuronal migration. Sorting of MGE-derived cortical INs out of the subpallium and their subsequent radial migration have been shown to rely on the coexpression of NRP1/2 which convey the chemorepellent force of SEMA3A/3F enriched in the striatal primordium and forming pallium (Marin and Rubenstein 2001; Nobrega-Pereira et al. 2008; Tamamaki et al. 2003). However, Plexin signaling is of high complexity and, in several cases, a single Semaphorin can interact with a variety of different Plexins. Indeed, as described for intracortical and sympathetic axons, different cell types might exhibit distinct functional coupling of Plexins and Neuropilins to convey the same Semaphorin signal (Waimey et al. 2008; Chen et al. 2000; Giger et al. 2000; Sahay et al. 2003). While alternate Plexin/Neuropilin complexes might cooperate in SEMA3A signal transduction, we find here that the genetic deletion of *Plxna4* does not influence the tangential migration of hINs, but specifically affects their laminar positioning in the cortex, without altering neither the distribution of PNs and MGE-derived INs nor the production of SEMA3A by these latter (data not shown). Interestingly, a recent publication reports an overmigration of superficial layer PNs into L1 when *Plxna2* is deleted in addition to *Plxna4* (Hatanaka et al. 2019). This work however does not report any mispositioning of GABAergic neurons in L1 and surroundings, where we find a substantial increase of hINs in *Plxna4* mutants. Although their observation is qualitative, the mechanisms through which an additional loss of *Plxna2* would compensate for the lack of *Plxna4* remains unclear. Given the complexity of Plexin signaling and the influence of their differential expression within a cell, we cannot exclude that *cis/trans* combinatorial mechanisms might be involved (Suto et al. 2007). Here we provide clear evidences that SEMA3A signal to hINs is *Plxna4*-dependent and most probably requires NRP1 as coreceptor. Supporting these elements, *in vivo* overexpression of SEMA3A does not alter hINs location in *Plxna4*^*-/-*^ brains, suggesting that other Plexins do not cover a central role in mediating the repulsive SEMA3A signal. Additionally, NRP1 blockade rescues SEMA3A-induced collapse of hINs *in vitro*, whereas NRP2 appeared less central in this complex. Finally, in line with a previous work suggesting that main cortical interneuron classes interact with each other for correct allocation (Liodis et al. 2007), we demonstrate that SEMA3A secreted by MGE-derived INs is required to control the laminar positioning of hINs. To do this, we engineered an *in vivo* model in which MGE-derived INs normally reach and settle into the cortex, but cease to express *Sema3a*. Using this approach, we manage to alter the laminar allocation of hINs, without affecting the positioning of MGE-derived INs nor cytoarchitecture of superficial layers PNs. Consistent with the loss of a SEMA3A repulsive signal in deep layers, we find that hINs redistribute across cortical layers, losing their preference to superficial ones. Similarly to previous work done on *Sema3a*^*-/-*^ mice (Andrews et al. 2017), we do not observe significant changes in the overall density of hINs in *Lhx6*^*Sema3a*-/-^ brains. This precluded the hypothesis that, due to the lack of SEMA3A, hINs misplaced in deep layers of *Lhx6*^*Sema3a*-/-^ brains could have bypassed cell death, a developmental event occurring during the two first postnatal weeks (Wong et al. 2018; Wong and Marin 2019) and shown to also be induced by lengthened exposure to SEMA3A (Bagnard et al. 2001; Ben-Zvi et al. 2008). Another observation of mention is that a small fraction of hINs (∼15%) locates in deep cortical layers in the control conditions and thus remains unsensitive to SEMA3A cell-extrinsic signaling. Interestingly, we find that a similar fraction of hINs expresses *Sema3a* mRNA, thus suggesting that *cis*-expression of SEMA3A could prevent the binding of PLXNA4 to *trans*-SEMA3A, a phenomenon already investigated in the sensory system between PLXNA4 and SEMA6A (Haklai-Topper et al. 2010). Whether other SEMAs may interact with *Plxna4*-expressing hINs is not precluded. Specifically, SEMA6A has been shown to signal with PLXNA4 in retinal, hippocampal, cortico-spinal tract and thalamocortical afferences development (Matsuoka et al. 2011; Mitsogiannis, Little, and Mitchell 2017; Runker et al. 2008; Suto et al. 2007). However, no mispositioning of neocortical INs has been reported in *Sema6a* knock-out mice, arguing against a pivotal role of SEMA6A in hINs lamination (Hatanaka et al. 2019). In conclusion, we propose a model in which the developing serotonergic system is triggering hINs to gradually express PLXNA4 as they migrate towards the CP. Concomitantly, MGE-derived INs, established in deep cortical layers, secrete SEMA3A and, by chemorepulsion, drive PLXNA4-expressing hINs to settle in superficial layers. A complementary attractive force driven by PNs is not precluded, but its nature is yet to be clarified. Overall, these results support a novel mechanism through which deep layer INs (i.e., MGE-derived) cell-extrinsically control the intracortical laminar positioning of their superficial layer counterparts (i.e., POA/CGE-derived).

Finally, the phenotypical discrepancies of hINs misplacement we observe among *Htr3a*^*-/-*^ (Murthy et al. 2014), *Plxna4*^-/-^ and *Lhx6*^*Sema3a*-/-^ models may be due to modifications occurring at different steps and on different targets of a single mechanism (Fig. 8). In hINs *Htr3a* modulation is shown to impact the regulation of several other genes that may be somehow involved in at least one of the several aspects of their migration and establishment in the developing cortex. In *Plxna4*^-/-^ brains, substitutive PLXN/NRP complexes could respond to other signaling cues or differently to SEMA3A, a phenomenon observed in other systems (Smolkin et al. 2018; Yaron et al. 2005). By contrast, *Lhx6*^*Sema3a*-/-^ brains are engineered to carry a selective deletion of SEMA3A in MGE-derived INs, thereby, keeping cell-intrinsic programs of hINs unaffected. Although beyond the scope of the present study, it would be necessary to assess maturation and integration into circuits of both misplaced hINs and SEMA3A-depleted MGE-derived INs in *Lhx6*^*Sema3a*-/-^ brains.

## Acknowledgments

We thank Valérie Castellani and Marc Tessier-Lavigne for providing *PlexinA4*-KO mice and Theofanis Karayannis for providing *Lhx6*-Cre mice. We thank Julien Prados and Joan Badia for help with databases mining, Nicolas Liaudet for Matlab script design and Wafae Adouan and Michèle Brunet for technical assistance. We would also thank members of Dayer and Jabaudon laboratories for fruitful discussions and insights. This work was supported by the Swiss National Foundation (SNF) grant (31003A_175445) and Synapsy grant (51NF40-185897) to A.D.

## Methods

### Animals

Animal experiments were approved by the Geneva animal care committee (authorizations GE/176/17 and GE/173/19) and performed according to the Swiss guidelines. Mice were housed in the conventional area of the animal facility of the University Medical Center, under controlled temperature (22±2°C) and dark/light cycles (12h each). Food and water were provided *ad libitum*. Timed pregnancies were obtained by overnight mating and the following morning counted as embryonic day (E) 0.5. In order to account for biological variability, at least two litters from different parents were sampled in each experiment. Both sexes were equally accounted for analyses. Transgenic and control animals were littermates, obtained by heterozygous mating (genotype was determined *a posteriori*). All animal lines were described previously: Tg(Htr3a-EGFP)DH30Gsat/Mmnc (*Htr3a*-GFP) (MMRRC 000273-UNC) (Lee et al. 2010; Murthy et al. 2014; Vucurovic et al. 2010), Tg(Gad2-EGFP)DJ31Gsat (*Gad65*-GFP) (MMRRC 011849-UNC) (Murthy et al. 2014; Lopez-Bendito et al. 2004), Htr3a^tm1Jul^ (*Htr3a*^-/-^) (Murthy et al. 2014; Zeitz 2002); Plxna4^tm1Matl^ (*Plxna4*^-/-^) (Yaron et al. 2005); B6;CBA-Tg(Lhx6-icre)1Kess/J (*Lhx6*^*cre*^) (JAX 026555) (Fogarty et al. 2007), *ICR*.*Cg-Sema3A*^*tm1*.*2Tyag*^*/TyagRbrc* (*Sema3A*^*fl/fl*^) (RBRC01106) (Taniguchi et al. 1997), Gt(Rosa)26Sor^tm9(CAG-tdTomato)Hze^/J (*Rosa26*-tdTOM^*fl/fl*^) (JAX 007909). CD1 mice (Wild-type) were purchased from Charles River Laboratory.

### Tissue preparation

Pregnant females were euthanized by intraperitoneal (i.p.) injection of thiopental (50mg/kg, Inresa), embryos exposed by caesarian cut and brains dissected out and fixed overnight (O.N.) at 4°C in 0.1M phosphate buffer pH7.4 with 4% paraformaldehyde (PFA). For postnatal ages, mice were anesthetized by i.p. injection of thiopental and transcardially perfused with 0.9% saline/liquemin followed by ice-cold PFA. Coronal brain sections were cut on a vibratome (Leica VT1000S) at 60µm for immunohistochemistry (IHC) or at 80-100µm for *in situ* hybridization (ISH). If not immediately processed, slices were stored at - 20°C in an ethylene glycol-based cryoprotective solution.

### Immunohistochemistry

For IHC, free-floating slices were rinsed in phosphate buffer saline (PBS) pH7.4 with 0.3% Triton-X100 and 0.1% Azide (PBST/azide) and blocked in PBST/azide with 2% Normal Horse Serum (NHS). Primary antibodies applied for two overnight at 4°C in the latter solution. The following primary antibodies were used: rat anti-CUX1 (1:250, Santa Cruz Biotechnology), chicken anti-GFP (1:2000, Abcam), goat anti-GFP (1:2000, Abcam), rabbit anti-GFP (1:500, Millipore), rabbit anti-Neuropilin1 (1:100, Abcam), goat anti-Neuropilin2 (1:50, Santa Cruz Biotechnology), goat anti-PROX1 (1:250, R&D System), mouse anti-Parvalbumin (1:2000, Swant), rabbit anti-PlexinA4 (1:500, Abcam), rabbit anti-Semaphorin3A (1:500, Abcam) rat anti-Somatostatin (1:500, Millipore), goat anti-tdTomato (1:1000, Sicgen). After washing the unbound primary antibodies and azide in PBST, secondary donkey Alexa-488, -546/555 and -647 antibodies (Thermo Fisher, Jackson) raised against the appropriated species were applied at 1:500 dilution in PBST for 1h at room temperature (RT). Sections were then rinsed in 0.1M phosphate buffer pH7.4 (PB), counterstained with Hoechst 33258 (1:5000, Invitrogen) and mounted with mowiol on gelatinized microscopy slides.

### In situ *hybridization*

The procedure was performed under RNase-free conditions and all solutions were added with 0.1% diethyl pyrocarbonate (DEPC, Sigma). Sections were quenched with 2% H_2_O_2_ and digested in 10µg/ml Proteinase K (BioLabs). The reaction was stopped with 2mg/ml Glycine (Sigma) and slices post-fixed in DEPC-PFA/2% glutaraldehyde (Sigma). The following plasmid probes were used: *Htr3a* (restriction: HindIII-HF, polymerization: T7; gift from B. Emerit), *Lhx6* (restriction: NotI, polymerization: T3; gift from M. Denaxa), *Sema3a* (restriction: NotI, polymerization: T7; gift from J.-P. Hornung), *Plexina4* (restriction: EcoRV, polymerization: T7; gift from V. Castellani). 5µg of the desired probe was applied to the sections and incubated O.N. at 65°C in a solution containing 50% deionized formamide (Sigma-Aldrich), 5x SSC pH7.0 (Invitrogen), 1% SDS (Sigma), 10mg/ml Salmon Sperm (Invitrogen) and 50µg/ml Heparin. Unbound probe was removed in formamide-based solutions with decreasing salts concentrations, slices blocked with 10% Blocking Reagent (Roche) in 100mM maleic acid/150mM NaCl/0.01% Tween20 pH7.5 and incubated O.N. at 4°C with alkaline phosphate anti-DIG (1:2000, Roche). For chromogenic revelation, slices were incubated with 4-Nitroblue tetrazolium chloride (NBT, Roche) and 5-bromo-4-chloro-3-indolyl-phosphate (BCIP, Roche) in NTMT pH9.5 solution (100mM NaCl, 100mM Tris-HCl, 50mM MgCl_2_, 0.1% Tween20) and mounted with Aquamount (Thermo Fisher Scientific). Fluorescent substrate was revealed with Fast Red tablets (Kem-En-Tech) dissolved in 0.1M Tris pH8.2. IHC and mounting were then performed as described above.

### Image acquisition and layering and density analyses

Images were acquired with an inverted confocal microscope (Nikon A1r), equipped with oil-immersion 40x (CFI Plan Fluor 40x/1.3) and 60x (CFI Plan Apo VC H 60x/1.4) objectives (Nikon). Bright-field images were acquired with a widefield microscope (Axioscan.Z1, Zeiss), equipped with 10x objective (Plan-Apochromat 10x/0.45, Zeiss). For illustration purposes only, images were slightly modified for gamma gain and brightness and applied despeckle filter with Photoshop (v. CC and 2020, Adobe Creator).

Cell quantifications (layering, density and colocalization) in the somatosensory cortex were achieved manually with cell counter plugin in Fiji (ImageJ). Layers and compartments of somatosensory cortices were defined according to Hoechst staining density. SEMA3A intensity analysis was carried on Fiji (ImageJ), by defining developing brain compartments in which mean fluorescence intensity was measured. Each value was normalized with the mean intensity of the corresponding white matter compartment.

### Acute slice preparation

For embryonic tissue, pregnant females were euthanized with thiopental, uterine horns exposed by caesarian cut and embryos heads collected in ice-cold Ca^2+^/Mg^2+^-free Hank’s balanced salt solution (HBSS, Gibco). For P0 tissue, pups were decapitated and heads collected in ice-cold HBSS. Brains were rapidly dissected out, and embedded in 3% low-melting point agarose (LMP-agarose, Roth). 250µm-thick slices at the level of the somatosensory cortex were then cut in ice-cold HBSS on a Vibratome (Leica VT1000S) and kept in their surrounding agarose. Slices were then placed in an incubator for recovery. Details concerning the age, orientation and handling of the slices are given in the corresponding following sections.

### FACS and Microarray

Acute coronal slices at E14.5, E18.5 and P2 from *Htr3a*^*+/+*^;*Gad65*-GFP and *Htr3a*^*-/-*^;*Gad65*-GFP mice were microdissected under a fluorescent scope (Leica M165FC). For each time-point and condition, triplicates from 3 different litters were obtained by pooling at least 4 brains. Tissue was dissociated with 0.25% Trypsin-EDTA (Life Technology) and pellet resuspended in Neurobasal medium (NBM, Gibco) supplemented with 2% B-27 (Gibco), 2mM 100x GlutaMAX (Gibco), 1% Penicillin-Streptomycin, 2mM N-Acetylcysteine and 1mM Sodium Pyruvate (i.e., supplemented NMB). GFP^+^ cells were isolated by fluorescence activated cell sorting (FACS) with FACS VantageSE and collected in RNA later (Invitrogen). RNA extraction was achieved with RNeasy Mini kit (Quiagen) according to manufacturer’s protocol and microarray were performed by the Genomic Facility at the University of Geneva as previously described (Murthy et al. 2014).

### In utero *electroporation*

Pregnant mice were anesthetized by inhalation of 2.5% isofluorane and O_2_ 2l/min and placed on a temperature-controlled pad. Under continuous maintenance of anesthesia, eyes were protected with gel drops (Viscotears) and analgesic (1µl of 0.5% Temgesic, Scherning-Plough) given subcutaneously. The shaved abdomen was sterilized with Betadine (MundiPharma), covered with a sterile defenestrated pad and uterine horns were exposed by caesarian cut. Approximately 700nl of either *pUb6*-tdTOM or *pUb6*-*Sema3a*-tdTOM plasmid (2µg/ml with 1% Fast Green, Sigma) was delivered into the lateral ventricle of embryo’s brain through a beveled glass pipette applied to a Picospritzer (Parker). Tweezers-type electrodes (CUY611P3-1, NepaGene) were placed on the brain at ∼35° to the horizontal plane, with the positive pole on the side of the injected ventricle and directed towards the dorsal pallium. Five electric pulses of 50V (50ms, 950ms interval) were applied with a squared wave electroporator (CUY21-SQ, NepaGene). Finally, horns were placed back in the abdominal cavity, muscles and skin were each stitched (Ethicon) and antibiotic (Fucidin 2%, Leo) applied on the scar. Mice were allowed to wake up and recover on a warm pad and embryos left to develop until the age of interest. Mothers’ health status was ensured every day until delivery and Paracetamol dissolved in water was given when needed (1g/l). Tissue collection, IHC, image acquisition, cell quantifications and analyses were performed as described above.

### Primary cell culture, peptide application and immunocytochemistry

CGEs from E14.5 *Plxna4*^*+/+*^;*Htr3a*-GFP or *PlxnA4*^-/-^;*Htr3a*-GFP coronal acute brain slices were microdissected under a fluorescence dissecting scope (Leica M165FC) and cells dissociated with 0.25% trypsin-EDTA (Life Technologies). Cells were resuspended in supplemented NBM and plated in homemade sterile polypropylene rings (6mm diameter) sealed with laboratory valve lubricant (Dow Corning) onto glass coverslips pre-coated with poly-D-lysin (Sigma) and laminin (Invitrogen). At DIV3, cells were treated for 1h at 37°C with 5nM of human SEMA3A Fc chimera (R&D System), dissolved in 1x Dulbecco’s PBS (DPBS, Gibco) with 0.1% bovine serum albumin (BSA, Sigma). In control conditions, only 1xDPBS/0.1%BSA (i.e., vehicle) was applied. To block ligand binding, cells were preincubated for 10min with 30µg rat anti-human neuropilin1 Fc chimera (R&D System) or 20µg goat anti-human neuropilin2 Fc chimera (R&D System) peptides before application of 5nM of SEMA3A as described before. Rings were fixed for 20min with pre-warmed 4% PFA and processed for immunocytochemistry (ICC). For ICC, cells were shortly permeabilized in 1x PBS with 0.1%Triton-X-100 and incubated for 1h30 in 10% NHS with chicken anti-GFP (1:2000, Abcam) and rabbit anti-PlexinA4 (1:500, Abcam) primary antibodies. Secondary donkey Alexa-488, -546/555 and -647 antibodies (Thermo Scientific, Jackson) raised against the appropriated species were applied (1:500) and finally cells were counterstained with Hoechst 33258 (1:5000) and mounted on gelatinized microscopy slides with mowiol. 20µm-thick stacks (2µm-stepped) were taken with an inverted confocal microscope (Nikon A1) equipped with an oil-immersion 60x objective (CFI Plan Apo VC H 60x/1.4, Nikon). Single-blind analysis of growth cone (GC) area was achieved with Fiji (ImageJ) using the maximal intensity projection (MIP) of the stacks. In order not to overrepresent one of the experimental replicates in a given condition, we performed equal random samplings of the quantified cones within the different conditions.

### Live tissue preparation, growth cones imaging and analysis

E17.5 *Plxna4*^*+/+*^;*Htr3a*-GFP and *PlxnA4*^-/-^;*Htr3a*-GFP acute coronal brain slices were cut, placed on floating nucleopore track-etched polycarbonate membranes (Whatman) and allowed to recover in supplemented NBM for at least 1h in an incubator (37°C, 5% CO_2_). Slices were then transferred and anchored in a superfusion chamber under an inverted confocal microscope (Nikon A1r), equipped with oil-immersion 60x objective (1.4 Plan Apo VC H, Nikon). 10µm-thick stacks of GFP^+^ GCs (1µm-stepped) were acquired with a 3x numerical zoom every 2min for 1h30 with resonant laser scanning at the level of the cortical plate. During the first 10min, sections were superfused with either vehicle (i.e., 0.1%BSA/1xDPBS) or 100nM human SEMA3A Fc chimera (R&D System) diluted in warm (37°C) oxygenated artificial cerebrospinal fluid (ACSF; 95% O_2_, 5% CO_2_) and finally with ACSF alone for the rest of the imaging session. Stacks were piled up to obtain MIP and time sequences aligned with StackReg plugin in Fiji (ImageJ). Area and GFP average intensity of GCs were calculated by contour tracing on Fiji (ImageJ). To capture at best the effect of SEMA3A on individual GC size and average intensity, both values were expressed as the percentage of the same value recorded at the beginning of the imaging. Paired statistical analyses were performed using the means of area and fluorescence intensity during the first 20min (Period 1, P1) compared to the last 20min (Period 2, P2) of the imaging session.

### HEK-T293 inverted drops, engraftment, imaging and analysis

HEK-T293 cells were maintained in culture flasks (Falcon) with Dulbecco’s Modified Eagle Medium (DMEM, Gibco) supplemented with 1% penicillin-streptomycin and 10% fetal calf serum (FCS, Gibco) in an incubator (37°C, 5% CO_2_). One fifth of the HEK-T293 culture was split into two petri dishes of 6cm diameter (Falcon) and transfected with lipofectamine (Invitrogen) and 8µg of either *pUb6*-tdTOM (i.e., control) or *pUb6*-*Sema3a*-tdTOM (i.e., SEMA3A overexpression) plasmid. Cells were then dissociated with 0.25% trypsin-EDTA (Life Technologies), pellet resuspended in supplemented DMEM and hanging drops placed on the lid of a humidified petri dish. Pseudotissue was allowed to form over two days in an incubator (37°C, 5% CO_2_). On the experimental day, hanging drops of transfected HEK-T293 cells were dissected in L-15 medium (Gibco) to obtain pieces of tissue of about 100×100µm. Dissected chunks were then engrafted in the cortical plate of E17.5 *PlxnA4*^*+/+*^;*Htr3a*-GFP acute coronal slices, placed on Millicell membranes (Millipore). Slices were allowed to recover on supplemented NBM in an incubator for at least 2h before imaging. Control and SEMA3A overexpression conditions were represented on each slice (one per hemisphere). Millicell containing the slices was then transferred to a supplemented NMB-filled Fluorodish (WPI) for time-lapse imaging. Acquisition was achieved with an inverted confocal microscope (Nikon A1r) equipped with long-working distance 20x objective (0.45 CFI ELWD Plan Fluor, Nikon). Microscope chamber was kept at 37°C, with 25l/h continuous flux of 5% CO_2_. 50µm stack (3µm-stepped) images were taken with resonant laser every 10min for 12h. Images were piled up to obtained MIP and sequences aligned with StackReg plugin on Fiji (ImageJ). Areas at 50µm and 100µm away from the patch were automatically defined with a Matlab-based script (developed by N. Liaudet, Bioimaging Facility) along 10h of movie (the first 2h were excluded to allow sections to settle in the imaging environment) and each hour, GFP^+^ cells in the given areas were manually counted with Fiji (ImageJ) and their density calculated. GFP^+^ cell movements into the HEK-T293 engraftment were counted and shown as event per 100µm of engraftment border along 10h.

### Ex vivo *slice engraftment, imaging and analysis*

Acute coronal sections from P0 *Plxna4*^*+/+*^ or *Plxna4*^*-/-*^;*Htr3a*-GFP brains were prepared as described above and SVZ/IZ stream microdissected in ice-cold L-15 medium (Gibco) under a fluorescence scope. Dissected tissue was then isotopically engrafted into isochronic wild-type acute coronal sections (previously left to recover on supplemented NBM in an incubator at 37°C, 5% CO_2_), placed on L-15 filled Millicell membranes. Millicell containing engrafted slices were kept for 24h on supplemented NBM in an incubator (37°C, 5% CO_2_), in order to allow GFP^+^ cells to start invade the host tissue. Live imaging was then started and carried along 12h as described above. Regions of interest were defined with help of phase contrast imaging. The SEMA3A zone (corresponding to L5/6) was estimated as the upper half and the IZ as the lower half of a region comprised between the upper edge of the engraftment and the lower edge of the CP. Cell movements were analyzed with MTrackJ plugin in Fiji (ImageJ). The percentage of time spent in a zone was calculated as the number of frames a GFP^+^ cell body was in a given zone divided by the total number of frames this cell was trackable.

### Statistical Analyses

Microarray data were analyzed with Partek Genomics suites (v. 6.6 beta) and R program. mRNA expression levels (ProbeSet) were compared with 2-way ANOVA, uncorrected p < 0.01.

All statistical analyses were performed with GraphPad Prism software (v. 7.0a and 8.0.2). Normality of the samples was assessed with D’Agostino-Pearson test and non-parametric tests used when criteria were not fulfilled. Statistical significance was set at α = 0.05.

Details for statistical tests used for each analysis, as well as samples size and significance levels are thoroughly described in the corresponding Figure legend.

## Supplementary Figure Legends

**Supplementary Figure 1. *Plxna4* mRNA expression in *Htr3a*-GFP**^**+**^ **interneurons (hINs) tangentially migrating and settled in upper cortical layers (Related to Fig. 1)**

(A) *In situ* hybridization showing that tangentially migrating hINs at E14.5 do not express *Plxna4* (left, orange cell contours). At P2, *Plxna4* is expressed in hINs (white cell contours) having reached upper cortical layers (L2-4). Relative intensity levels indicate gradually increasing expression of *Plxna4* in hINs located in upper cortical layers (L2-4) at P2 (bottom right) (n = 2 brains, 27 cells in SVZ/IZ, 117 cells in L5/6, 51 cells in L2/4, 24 cells in L1). Scale bar: 50 µm (low magnifications), 25µm (A, high magnification), 10 µm (B, high magnifications).

**Supplementary Figure 2. Upper layers cytoarchitecture, migratory streams and molecular identity are preserved in *Plxna4***^**-/-**^ **brains; *Lhx6***^***cre***^ **mouse model validation (Related to Fig. 2)**

(A) Normal distribution of upper layer CUX1^+^ projection neurons in the somatosensory cortex of *Plxna4*^+/+^ (left) and *Plxna4*^-/-^ (right) brains at P21. Graph of mean ± SEM showing the ratio between the area occupied by CUX1 staining and the whole neocortical area. The two conditions seem to be comparable (*Plxna4*^+/+^: n = 6 sections from different brains; *Plxna4*^-/-^: n = 2 sections). Scale bar: 100µm.

(B) Tangentially migrating hINs from *Plxna4*^+/+^ (left) and *Plxna4*^-/-^ (right) brains at E14.5. No change in the density of hINs in the marginal zone (MZ) and intermediate/subventricular zone (IZ/SVZ) streams is observed in *Plxna4*^-/-^ brains (*Plxna4*^+/+^: n = 3 brains, 1850 cells; *Plxna4*^-/-^ : n = 3 brains, 2178 cells; n.s.: p = 0.3904; 2-way ANOVA). Scale bar: 100µm.

(C) Expression of the transcription factor PROX1 in hINs from *Plxna4*^+/+^ (left) and *Plxna4*^-/-^ (right) brains at P21 in the somatosensory cortex. The fraction of hINs expressing PROX1 is similar in *Plxna4*^*-/-*^ as compared to *Plxna4*^+/+^ brains (*Plxna4*^*+/+*^: n = 5 brains, 2451 cells; *Plxna4*^*-/-*^: n = 3 brains, 1356 cells; n.s.: p = 0.5714; Mann-Whitney test). Scale bar: 100µm.

(D) Coronal section of P0 *Lhx6*-tdTOM brain showing *Lhx6* mRNA expression (left), but not *Htr3a* mRNA (right) in recombined cells. Scale bar: 100µm (low magnification), 20µm (high magnification).

(E) Coronal section of P21 *Lhx6*-tdTOM brain showing immunostaining against Parvalbumin (PV, left) and Somatostatin (SST, right), common markers for MGE-derived INs. Analysis shows that virtually all recombined tdTOM^+^ cells express either PV or SST (n = 2 brains, 1522 cells). Scale bar: 200µm.

(F) ISH on coronal section of *Htr3a*-GFP brain at P0, showing that *Htr3a*-GFP^+^ INs (hINs) only rarely express *Sema3a* mRNA (11.54 ± 1.62 %; n = 3 brains, 1227 cells). Scale bar: 100µm (low magnification), 50µm (high magnification).

All data are mean ± SEM.

VZ: ventricular zone, CP: cortical plate

**Supplementary Figure 3. Growth cone (GC) areas of dissociated *Htr3a*-GFP**^**+**^ **interneurons (hINs) after Semaphorin3A (SEMA3A) application (Related to Fig. 3)**

(A) Graphs showing GC areas of *Plxna4*^+/+^ (white) or *Plxna4*^-/-^ (grey) hINs after vehicle or SEMA3A application (*Plxna4*^+/+^ vehicle: n = 3 experiments, 63 GCs; SEMA3A: n = 7 experiments, 427 GCs; ****p < 0.0001; *Plxna4*^-/-^ vehicle: n = 5 experiments, 105 GCs; SEMA3A: n = 5 experiments, 365 GCs; n.s.: p = 0.4326). Statistics are done with Kruskal-Wallis test with Dunn’s post-test.

(B) Graphs showing GC areas of *Plxna4*^+/+^ after SEMA3A application with NRP1 or NRP2 blocking peptides (NRP1: n = 4 experiments, 176 GCs; NRP2: n = 4 experiments, 244 GCs; n.s.: p > 0.99 and ****p < 0.0001, respectively).

Data are mean ± SEM

**Supplementary Figure 4. GFP fluorescence intensity in *Htr3a*-GFP**^**+**^ **interneurons (hINs) growth cones (GCs) and hINs density surrounding transfected HEK-T293 cells explants (Related to Fig. 4)**

(A) Graphs of means ± SD showing that at the beginning of the imaging session GC sizes and intensities are comparable between *Plxna4*^*+/+*^ hINs and *Plxna4*^*-/-*^ hINs in all experimental conditions (n.s.: p = 0.7565, 1-way ANOVA; n.s.: p = 0.0845,Welch’s ANOVA, respectively).

(B) GFP mean intensities (left) are measured every 2 min along a timeline consisting of 10 min vehicle (black) or SEMA3A (magenta) application, followed by 72 min of ACSF superfusion. Graph (percentage of the initial value) showing that GFP fluorescence of GCs of *Plxna4*^*+/+*^ hINs (n = 12; black open circles) is relatively constant along vehicle perfusion, but increases within an hour following SEMA3A application (n = 11; magenta open circles). GCs of *Plxna4*^*-/-*^ hINs are not affected by SEMA3A (n = 11; magenta full circles). Graphs of paired values (percentage of the initial value, right) show that GCs of *Plxna4*^*+/+*^ hINs increase in average GFP intensity (***p = 0.0007; paired t-test) from the initial period (P1; 0-20 min) to the final period (P2; 60-80 min) after SEMA3A application. Vehicle superfusion does not significantly impact GCs of *Plxna4*^*+/+*^ hINs (n.s.: p = 0.46; paired t-tests). GCs of *Plxna4*^*-/-*^ hINs are not affected by SEMA3A (n.s.: p = 0.061; paired t-tests).

(C) Graphs of mean ± SEM showing hINs density in the 50µm (left) and 100µm (right) areas (n = 5 explants for each condition, control explant: 9244 cells; SEMA3A explant: 5834 cells; 50µm area: *p < 0.01, **p < 0.001; 100µm area: n.s.: p > 0.05; 2-way ANOVA with Geisser Greenhouse correction and Tukey’s post-tests).

**Supplementary Figure 5. The laminar positioning of projection neurons is not affected by SEMA3A overexpression (Related to Fig. 5)**

(A) Illustrative images showing the distribution of superficial layer projection neurons (PNs) in the P21 somatosensory cortices of *Plxna4*^+/+^;*Htr3a*-GFP brains electroporated at E15 with *pUb6*-tdTOM (left, control) or *pUB6*-*Sema3a*-tdTOM (right) plasmid construct. Layering analysis shows similar laminar distribution of PNs in the control and SEMA3A overexpression conditions (control: n = 5 brains, 4293 cells; SEMA3A: n = 4 brains, 2419 cells; n.s.: p = 0.3677; 2-way ANOVA). Scale bars: 200µm.

**Supplementary Figure 6. PLXNA4 is required for *Htr3a*-GFP**^**+**^ **interneurons (hINs) to sense endogenous source of SEMA3A. Upper layers cytoarchitecture is not affected in *Lhx6***^***Sema3a*-/-**^ **brains (Related to Fig. 6)**

(A) Schematic showing the *ex vivo* engraftment paradigm. At P0, the SVZ stream of either *Plxna4*^*+/+*^ or *Plxna4*^*-/-*^;*Htr3a*-GFP brains was dissected and grafted isotopically into isochronic wild-type tissue. Time lapse imaging was performed over 12h after 1 day *in vitro* and speed and time spent in the different parts of the nascent neocortex analyzed.

(B) Graph showing average speed of engrafted *Plxna4*^*+/+*^ or *Plxna4*^*-/-*^;*Htr3a*-GFP^+^ interneurons (hINs) in IZ and SEMA3A-expressing zone (*Plxna4*^*+/+*^: n = 15 experiments, 223 cells; *Plxna4*^*-/-*^: n = 12 experiments, 189 cells; n.s.: p = 0.1048, ***p = 0.0007, 2-way ANOVA with Bonferroni’s post-test)

(C) Graphs showing percentage of time hINs spend in either IZ or SEMA3A-expressing zone (**p = 0.0079; ****p < 0.0001; 2-way ANOVA with Bonferroni’s post-test).

(D) Normal distribution of upper layer CUX1^+^ projection neurons in the somatosensory cortex of *Lhx6*^*Sema3a*+/+^ (left) and *Lhx6*^*Sema3a*-/-^ (right) brains at P21. Graph showing the ratio between the area occupied by CUX1 staining and the whole neocortical area. No differences are observed between the two conditions (n = 4 brains per condition; n.s.: p = 0.8857, Mann-Whitney test). Scale bar: 100µm.

Data are mean ± SEM.

