## Supplementary Figures for "PlexinA4-Semaphorin3A mediated crosstalk between main cortical interneuron classes is required for superficial interneurons lamination"

### SUPPLEMENTARY FIGURE 1

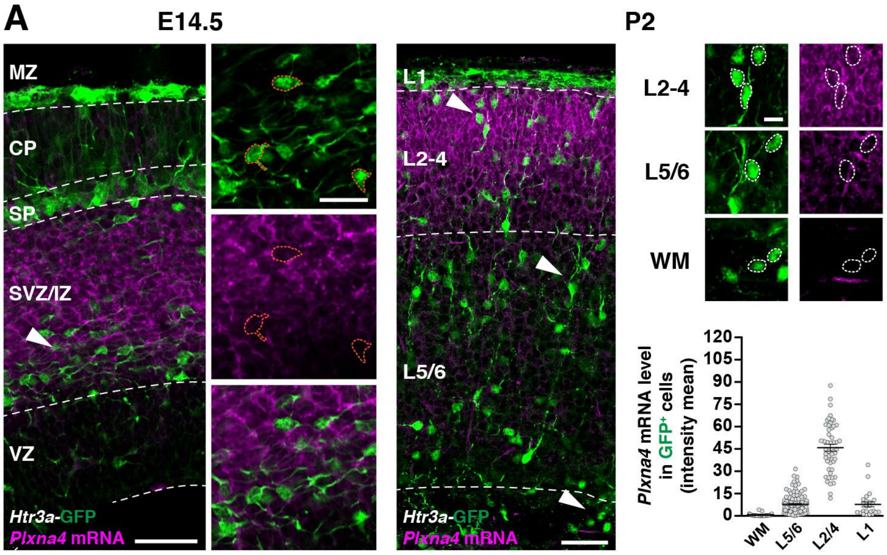

### SUPPLEMENTARY FIGURE 2

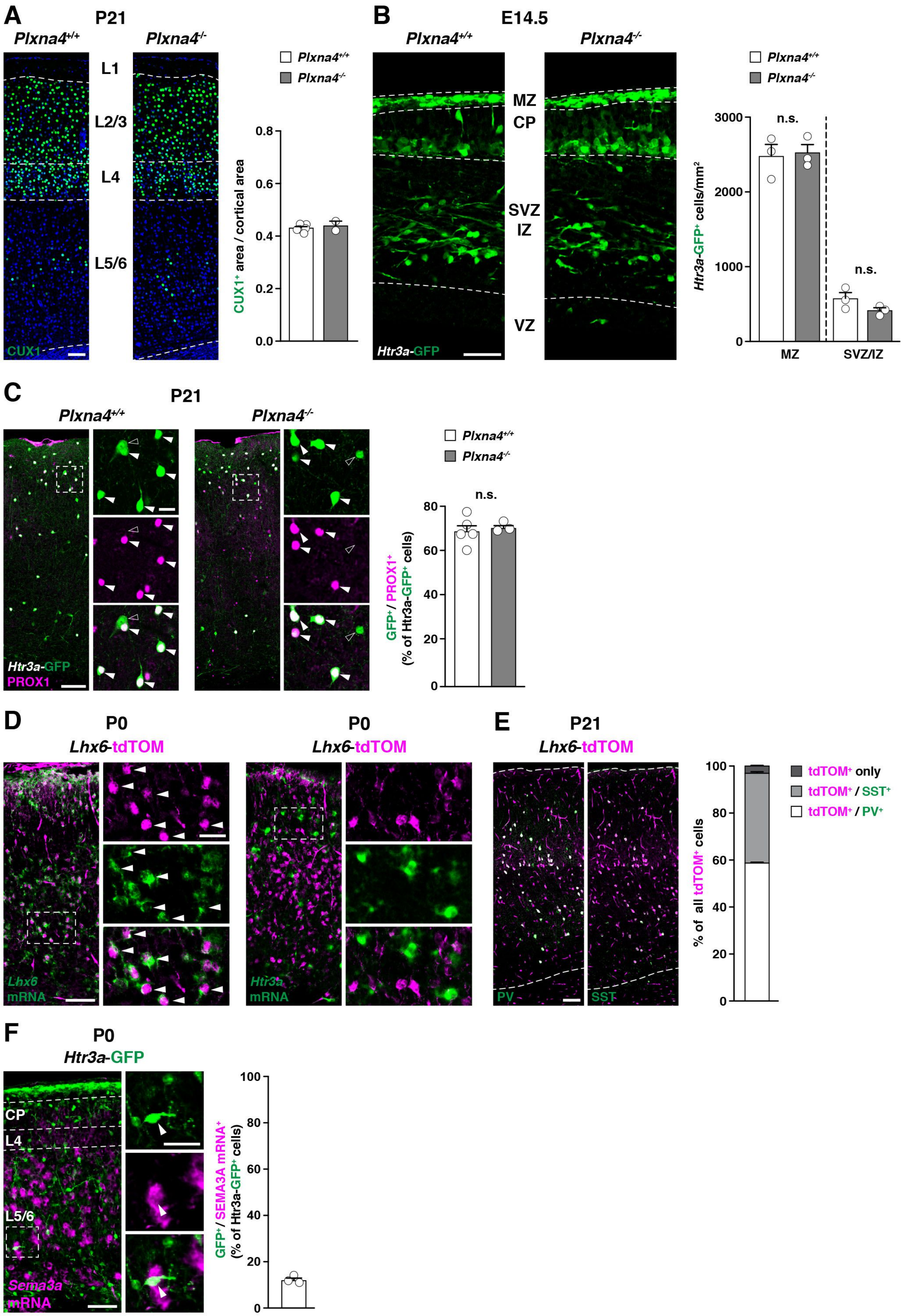

### SUPPLEMENTARY FIGURE 3

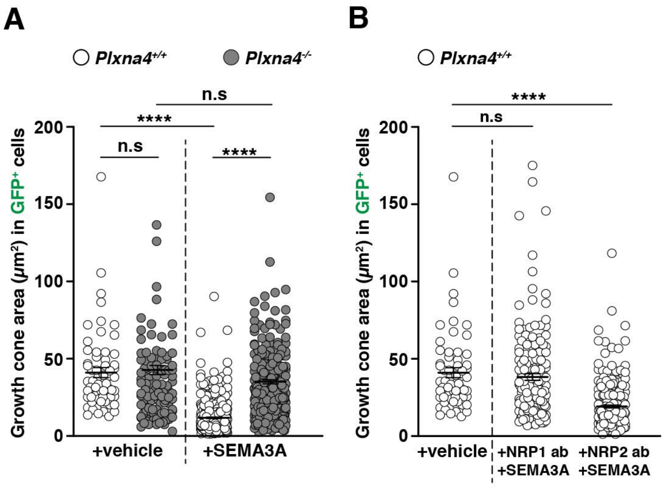

### SUPPLEMENTARY FIGURE 4

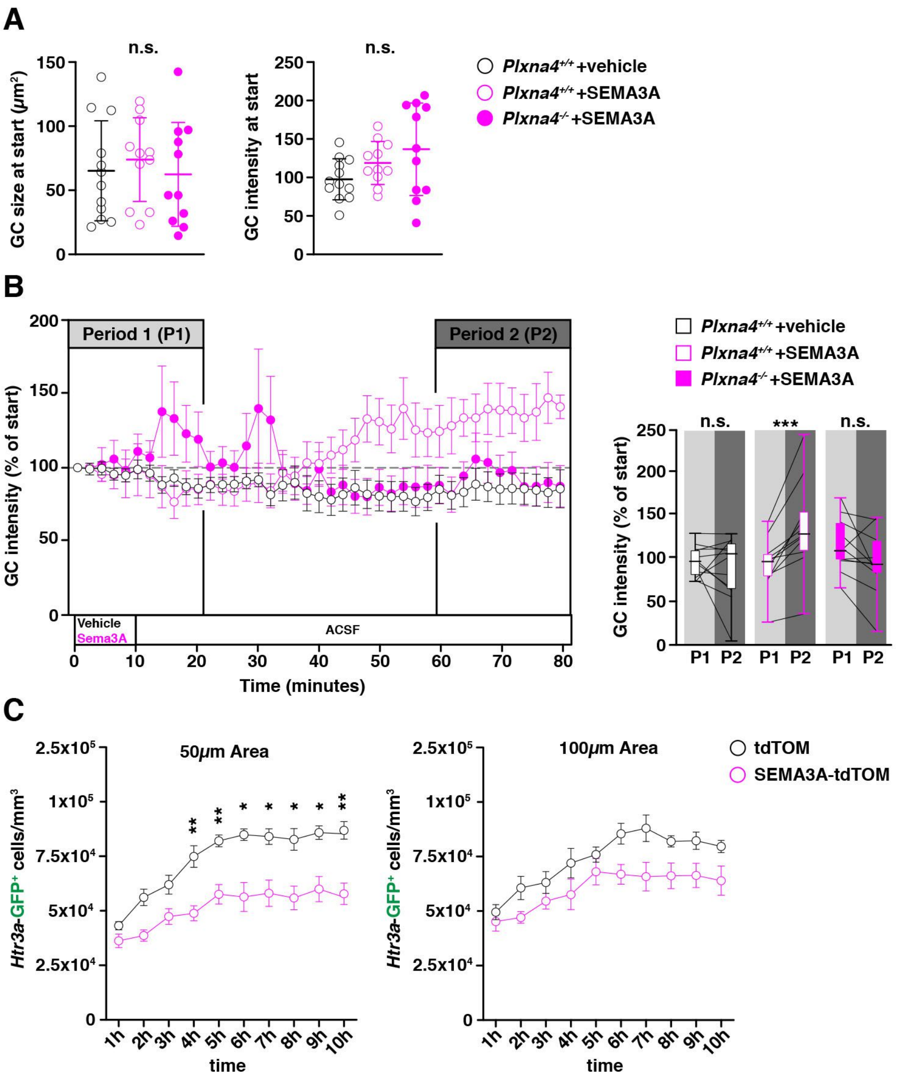

### SUPPLEMENTARY FIGURE 5

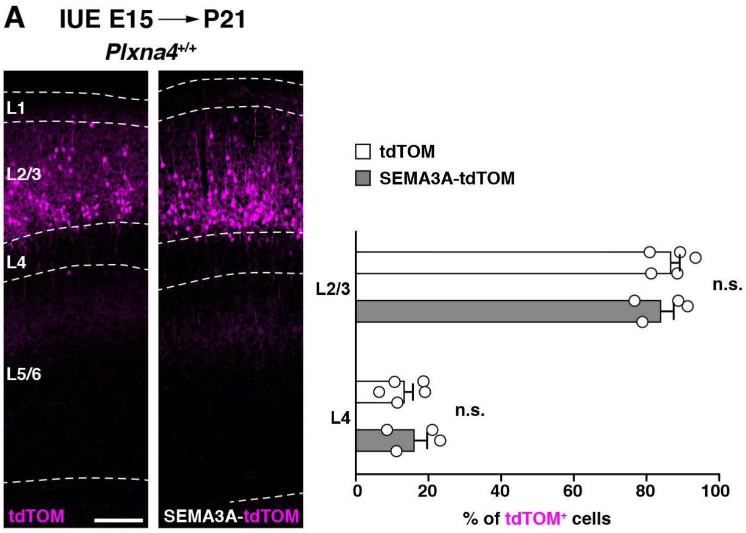

### SUPPLEMENTARY FIGURE 6

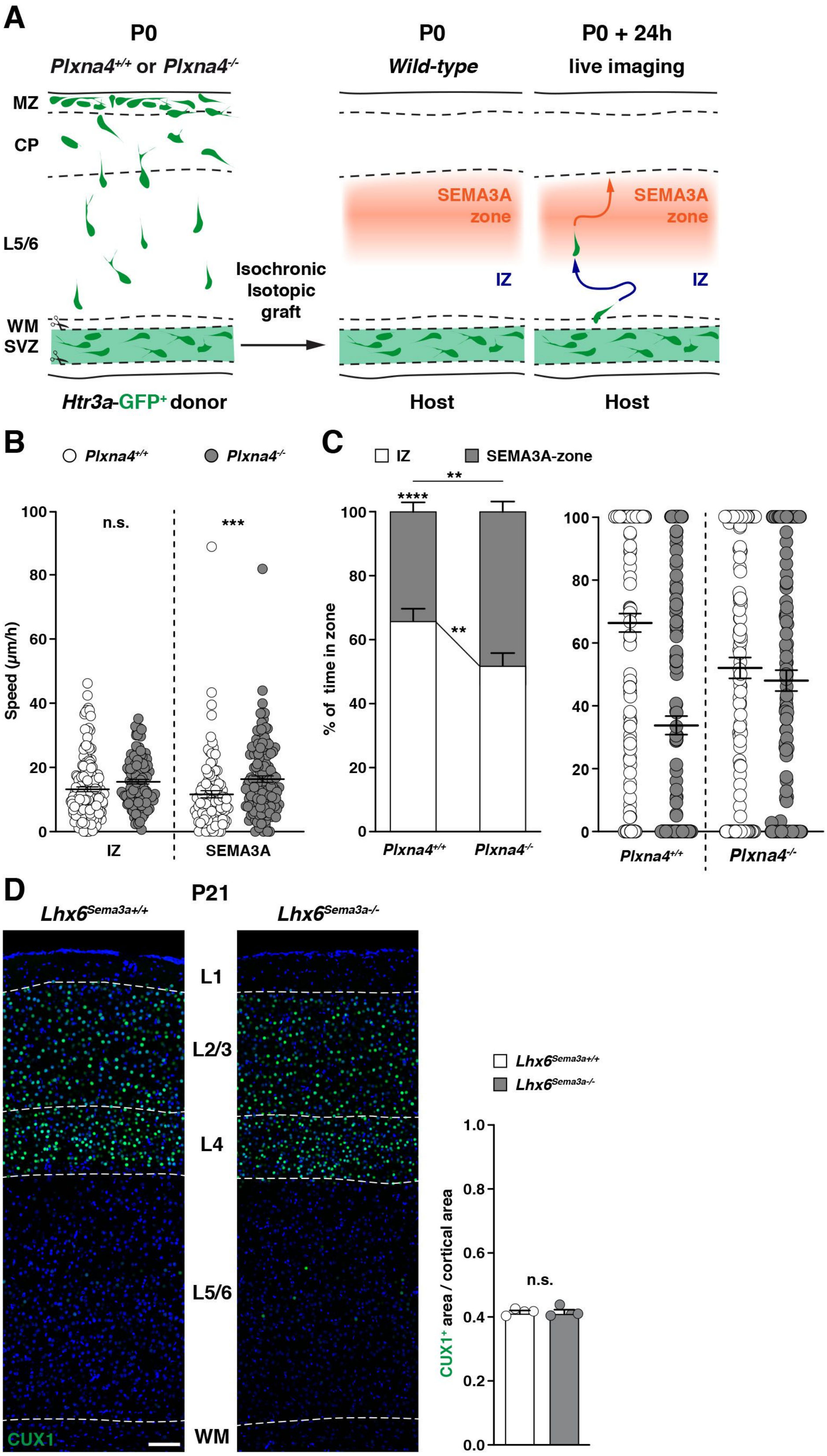
